# Systematic Identification of Post-Transcriptional Regulatory Modules

**DOI:** 10.1101/2023.02.27.530345

**Authors:** Matvei Khoroshkin, Andrey Buyan, Martin Dodel, Albertas Navickas, Johnny Yu, Fathima Trejo, Anthony Doty, Rithvik Baratam, Shaopu Zhou, Tanvi Joshi, Kristle Garcia, Benedict Choi, Sohit Miglani, Vishvak Subramanyam, Hailey Modi, Daniel Markett, M. Ryan Corces, Ivan V. Kulakovskiy, Faraz Mardakheh, Hani Goodarzi

## Abstract

In our cells, a limited number of RNA binding proteins (RBPs) are responsible for all aspects of RNA metabolism across the entire transcriptome. To accomplish this, RBPs form regulatory units that act on specific target regulons. However, the landscape of RBP combinatorial interactions remains poorly explored. Here, we performed a systematic annotation of RBP combinatorial interactions via multimodal data integration. We built a large-scale map of RBP protein neighborhoods by generating *in vivo* proximity-dependent biotinylation datasets of 50 human RBPs. In parallel, we used CRISPR interference with single-cell readout to capture transcriptomic changes upon RBP knockdowns. By combining these physical and functional interaction readouts, along with the atlas of RBP mRNA targets from eCLIP assays, we generated an integrated map of functional RBP interactions. We then used this map to match RBPs to their context-specific functions and validated the predicted functions biochemically for four RBPs. This study highlights the previously underappreciated scale of the inter-RBP interactions, be it genetic or physical, and is a first step towards a more comprehensive understanding of post-transcriptional regulatory processes and their underlying molecular grammar.

## INTRODUCTION

RNA binding proteins (RBPs) are crucial for regulating all stages of post-transcriptional regulation, from RNA splicing and nuclear export to translation and decay. Despite the limited number of RBPs encoded in the human genome (less than 1500) (Gerstberger, Hafner, and Tuschl 2014), they shephard more than 100,000 transcripts throughout their life cycles. The conventional “one RBP-one function-one regulon” model is therefore insufficient in capturing the multifaceted roles of RBPs (Glisovic et al. 2008). Instead, a more precise model is the “one RBP-many functions-many regulons” approach, which acknowledges that RBPs have multiple, context-dependent functions and regulate independent regulons. For instance, muscleblind-like 1 (MBNL1) regulates splicing when binding pre-mRNAs in the nucleus (Lin et al. 2006); while modulating RNA stability by binding 3’UTRs of target mRNAs in the cytoplasm (Fish et al. 2016). Similarly, SRSF1 simultaneously controls RNA processing in the nucleus and mRNA translation and decay in the cytoplasm (Twyffels, Gueydan, and Kruys 2011). These token examples illustrate the importance of recognizing the multifunctionality of RBPs in post-transcriptional regulation.

Although systems biology has been effective in providing tools to annotate RBPs, there is still a need for more research to characterize the context of these proteins’ regulatory functions. Several recent large-scale projects have focused on mapping different types of interactions between RBPs and their binding partners. For example, the ENCODE consortium has mapped over 120 RBP binding sites in the human transcriptome, while other projects have mapped the subcellular localization of hundreds of RBPs and RNAs, and investigated changes in gene expression profiles following protein knockdowns (Replogle et al. 2022; Fazal et al. 2019; Youn et al. 2018). As a result, the data characterizing interactions between RBPs and RNAs is available for many RNA binding proteins. However, as showcased above, not every RBP-RNA interaction has a similar impact on the fate of the target RNA (Corley, Burns, and Yeo 2020). The functional consequences of a given RBP-RNA interaction emerge from its molecular context, specifically through interactions with other RBPs that act on the same target. To capture post-transcriptional regulatory networks, we need a comprehensive map of multi-component RBP modules that effectively separate the set of bound RNAs into functionally distinct regulons.

In this study, we tackled this challenge by providing a systematic framework for mapping the RBP regulatory modules that form the core of these post-transcription regulatory networks. The challenge of this problem arises from the fact that functional interactions between RBPs are more complex than simple physical interactions. While RBPs can co-localize or directly interact, they also often cooperate by binding the same RNAs simultaneously or sequentially. Moreover, as RBPs form regulatory pathways, we also expect genetic interactions between RBPs, which manifest themselves as epistatic interactions in phenotypic measurements. Since no single biochemical assay is capable of capturing such a variety of interaction modes, a multi-modal approach is required to reliably annotate these RBP-RBP interactions. To capture these, we examined three forms of functional RBP-RBP entanglements: (i) two RBPs are co-localized in close physical proximity to each other, likely as part of a complex, (ii) two RBPs bind the same RNA targets, either simultaneously or at different times and locations, and (iii) two RBPs are part of the same pathway, and their modulations result in similar downstream transcriptomic changes. By combining data from these three independent modalities within a unified statistical and analytical framework, we have successfully captured RBP regulatory modules in their most generalizable form.

We further demonstrated that functional RBP interaction networks are significantly more complex than previously appreciated. We also used our multi-modal integrative map of RBP interactions to match individual RBPs to their functional regulons and revealed many previously unknown post-transcriptional regulatory processes governed by combinatorial interactions between RBPs. We have showcased and validated multiple instances of these molecular mechanisms using genetic, molecular, and biochemical approaches. Our discovery of a large repertoire of functional interactions between RBPs, many of which were previously unknown, provides a rich resource for the community to better study the role of post-transcriptional regulatory programs in health and disease.

## RESULTS

### Multi-modal approaches for capturing RBP-mediated interactions

In order to cover multitudes of functions for thousands of distinct target regulons, RBPs assemble into units of post-transcriptional control in a combinatorial manner (Cho et al. 2022). This process can be abstracted as a 4-layer network (Fig. 1A), where the first layer represents RBPs and the final layer their functionally distinct target regulons. In the intermediate layers, RBPs that play related roles and broadly share protein partners, targets, or context, come together to form large groups that we term “regulatory modules”. Then, RBPs within each module assemble into smaller regulatory units, each carrying out a specific regulating function and targeting a defined regulon. The resulting combinatorial RBP interaction network allows a limited number of RBPs to fulfill diverse functional roles in post-transcriptional regulation that is required to govern all aspects of the life cycle for all RNAs in the cell.

**Figure 1.**
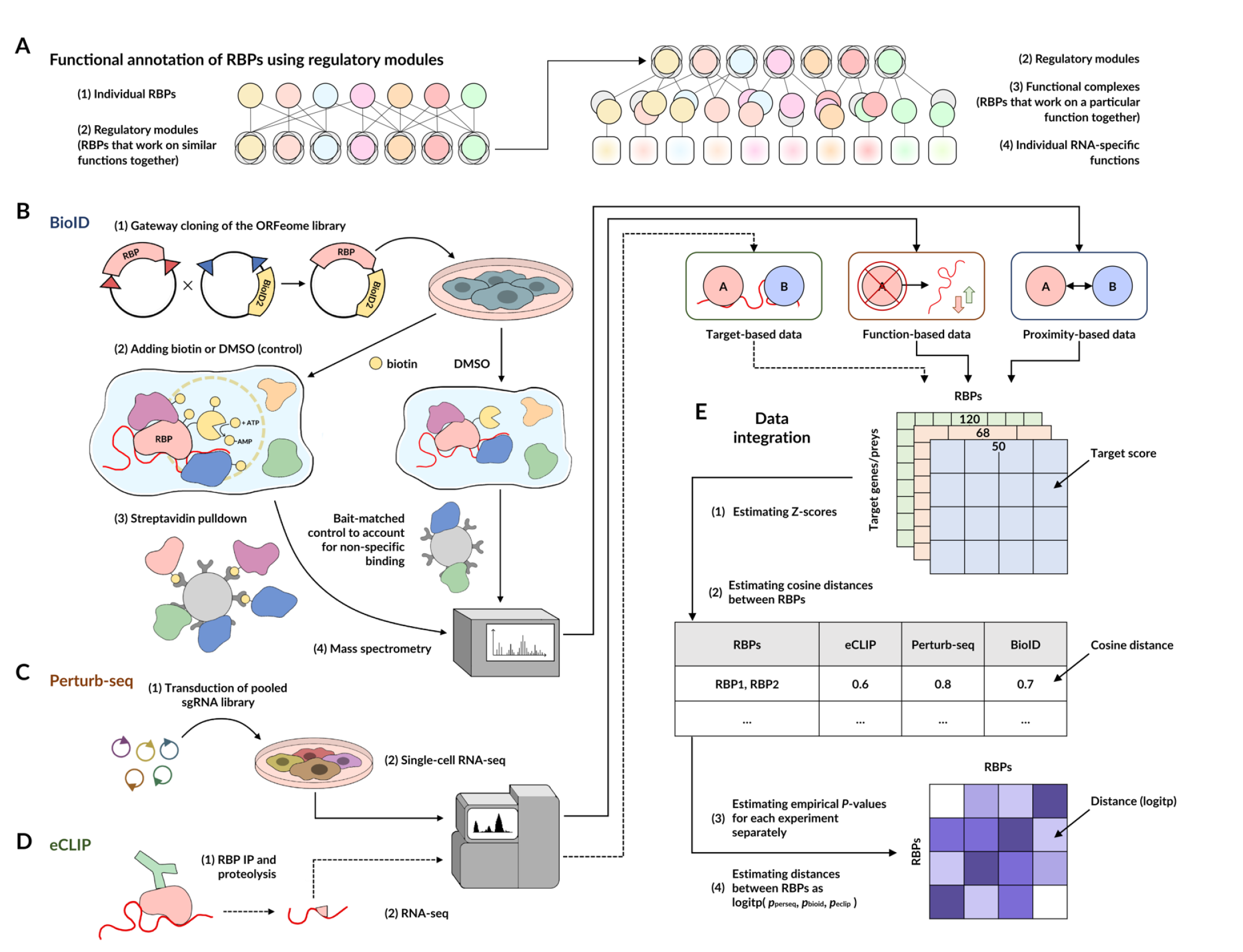
Workflow overview: generating an integrated regulatory interaction map of RNA-binding proteins. **(A)** Framework for functional annotation of RBPs. Layers 1-2: individual RBPs assemble into large groups, termed “regulatory modules”. Layer 3: functional complexes consist of RBPs, recruited from the same regulatory module. Layer 4: a functional complex implements a particular functional role on a defined group of RNA targets. **(B-D)** The results of BioID2, Perturb-seq and publicly available ENCODE eCLIP assays that were independently processed and normalized across RBPs **(E)**. The resulting Z-scores were used to estimate the cosine distance between all pairs of the tested RBPs and to calculate empirical p-values for RBP-RBP similarities. For each pair of RBPs, the p-values from three assays were aggregated as in (Mudholkar, George, and ROCHESTER UNIV NY DEPT OF STATISTICS 1977) to obtain a single measure of similarity between RBPs across the feature spaces from the three modalities. The resulting matrix of pairwise similarities was defined as an Integrated Regulatory Interaction Map that simultaneously captures physical and functional interactions between RBPs.

In order to broadly and systematically annotate regulatory interactions between RBPs, we combined data from three independent modalities, namely (i) BioID2-mediated proximity protein labeling, (ii) RBP Perturb-seq, and (iii) eCLIP RNA ENCODE dataset (Fig. 1, Fig. S1). First, to capture the protein neighborhood of each RBP, we used BioID labeling-based pulldown followed by mass spectrometry (D. I. Kim et al. 2016). We fused BioID2 to 50 RBPs (see Methods), and generated stable cell lines expressing each fusion construct in a human leukemia cell line K562. We then performed, in biological triplicates, streptavidin pull-down experiments followed by mass spectrometry. For each RBP, we also included matched controls by processing the same lines without a biotin pulse. This allowed us to generate a high-quality protein neighborhood dataset and to systematically identify co-localized RBPs. Second, we aimed to capture combinations of RBPs whose perturbations similarly impact the gene expression landscape of the cell. For this, we took advantage of Perturb-seq, a parallelized loss-of-function screen with rich single-cell transcriptomic readouts (Norman et al. 2019). We obtained transcriptome-wide gene expression measurements for 68 RBPs (representing a variety of regulatory processes; see Methods) knockdowns using single-cell RNA sequencing and used the resulting high-dimensional data to systematically delineate genetic interactions between RBPs (Norman et al. 2019). Finally, we re-analyzed the ENCODE eCLIP data to evaluate the extent to which pairs of RBPs bind to common RNA targets (Van Nostrand et al. 2016).

### Integrated RBP interaction maps to reveal regulatory modules

The abovementioned data modalities capture complementary aspects of regulatory interactions between RBPs. Therefore, integrating these sources of information is a critical step toward generating a more comprehensive and generalizable map of regulatory interactions (Fig 1, Fig. S1). To accomplish this, we first generated RBP-target interaction maps for each individual modality, where ‘target’ can be either neighboring protein (BioID), downstream gene (Perturb-seq), or target RNA (eCLIP; Fig. S2A-C). In order to make the values comparable between datasets, we z-score transformed them across the target features. RBPs that fall close to each other in this feature space, which reflects physical and functional proximity, likely function as part of the same regulatory modules. Therefore, for each data modality, we estimated pairwise cosine distances between RBPs and transformed them into empirical p-values to achieve a uniform scale for pairwise similarity. Finally, the three separately calculated p-values for RBP-RBP similarities (i.e. from BioID, Perturb-seq, and eCLIP respectively) were combined into a single unified probability score (Fig. S1) expressing the overall likelihood of functional interactions between pairs of RBPs.

The resulting ‘Integrated RBP Regulatory Map’, combining physical and genetic interactions, provides the means to elucidate the combinatorial regulatory logic underlying RBP-mediated post-transcriptional control of gene expression (Fig. 2A). Interaction maps are often interpreted by identifying proteins that cluster together into functional complexes in an unsupervised manner. And as shown in Fig. 2A, our map similarly captures a number of canonical RBP modules involved in key post-transcriptional regulatory programs such as ‘cytoplasmic translation’ and ‘mRNA splicing’. However, as stated above, RBPs carry out multiple independent functions, which are not captured by this “one RBP-one cluster” scheme. This is clearly highlighted by the many significant off-diagonal interactions that are observed in our clustered RBP map. And more importantly, our Integrated RBP Regulatory Map captures substantially more regulatory modules than an analogous map built based on the current state-of-the-art protein-protein interaction database, STRING-DB (Szklarczyk et al. 2018) (Fig. S2D).

**Figure 2.**
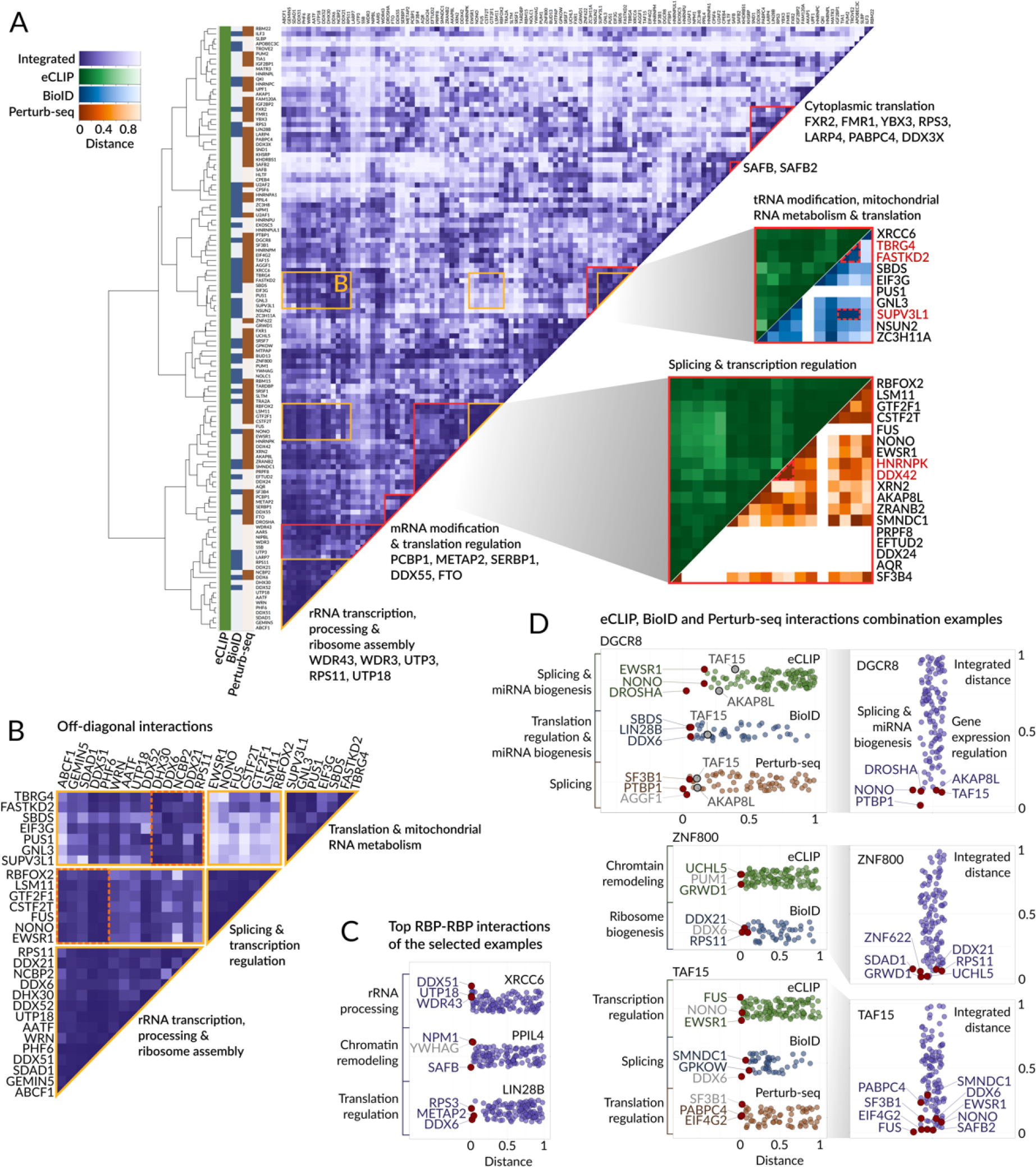
Post-transcriptional regulatory modules revealed by integrative analysis of RBP-RBP interactions. **(A)** Integrated RBP Regulatory Map: the heatmap of integrated distances between RBPs combining multiple sources of information for each RBP is shown in the middle. Each row and column represents an RBP, the integrated distances between RBPs are shown by color. RBPs are clustered based on their integrated distance using hierarchical clustering; the dendrogram is shown on the left. The presence of RBPs in the three source datasets is shown on the left with a colored map (green: eCLIP, blue: BioID, brown: Perturb-seq). Examples of known regulatory modules are highlighted in red and labeled; the RBPs with known functions that contribute to the module’s annotation are listed. For two exemplary regulatory modules, zoomed-in heatmaps are included. These selected heatmaps deconvolve the integrated distance into the distances between RBPs derived from individual datasets. Different color schemes were used for different source datasets: blue for BioID, orange for Perturb-seq, and green for eCLIP. The proteins discussed in the text are highlighted in red. The exemplary combinatorial interactions between regulatory modules are marked in yellow and shown separately in (B). **(B)** A slice of the Integrated RBP Regulatory Map showing combinatorial interactions between key regulatory modules. All the areas that are highlighted in yellow in (A) are shown. The individual regulatory modules are annotated. Two example associations are highlighted with orange dashed boxes: (1) association of the rRNA maturation factors ABCF1, GEMIN5, SDAD1, DDX51, and PHF6 with splicing machinery and (2) association of DHX30, DDX6, NCBP2, DDX21, and RPS11 and the regulators of translation and mitochondrial RNA metabolism. **(C)** Swarm plots showing the RBP partners of XRCC6 (top), PPIL4 (middle) and LIN28B (bottom). Each point represents a single RBP; the points are sorted based on the integrated distance of the given RBP to the query RBP. Each swarm plot is annotated with the common function of the lowest distance interacting partners. The three RBPs with the lowest distance (three top interacting partners) are labeled. Among them, those annotated with the common function are labeled in purple, and the rest are labeled with gray. **(D)** Swarm plots showing the RBP partners of DGCR8 (top), ZNF800 (center) and TAF15 (bottom) based on eCLIP, BioID, and Perturb-seq. Right: swarm plots showing the RBP partners of query RBPs based on the integrated distance, as in (C). The top interacting RBPs identified by the integrated or individual distances are labeled. Left: swarm plots showing the RBP partners of query RBP based on the individual source datasets. The swarm plots corresponding to individual datasets are color-coded in the same way as in (A). Each swarm plot is annotated with the common function of the lowest distance interacting partners and the three RBPs with the lowest distance are labeled as in (C). For DGCR8, two RBP partners, TAF15 and AKAP8L, are additionally highlighted in gray and labeled on the left, representing the effect of the distance integration procedure that recapitulates known DGCR8 involvement in the regulation of gene expression discussed in the text.

Our integrative approach brings together RBPs that form key regulatory modules and broadly recapitulates what is known about the functions of these RBPs; however, tracking the source of the signal to the individual input modalities is also often informative and can highlight the higher resolution of our integrative approach. For example, among the group of 18 RBPs that are collectively associated with splicing-related functions, and bind overlapping RNA targets based on the eCLIP data, HNRNPK and DDX42 are also associated with the regulation of p53-mediated apoptosis based on Perturb-seq results; and consistently, they have been previously shown to be upregulated in p53 mutant cells (Fig. 2A) (Escobar-Hoyos et al. 2020). Another example is the group of RBPs associated with mitochondrial and cytoplasmic translation. While eCLIP data brings together the rRNA-binding RBPs, proximity labeling data clearly distinguishes mitochondrial RBPs (TBRG4, FASTKD2, and SUPV3L1) from others (Fig. 2A).

RBPs form more functional interactions than captured by traditional PPI methods. We tested this hypothesis by comparing the interaction map built using protein-protein interaction data from STRING-DB database with the map built using our multi-modal integrated data from the current study. The integrated map, generally corresponding to the STRING-DB, showed larger regulatory modules, demonstrating the utility of a multi-modal data integration approach in revealing more functional interactions (Fig S2 D,E).

### Combinatorial interactions between RBPs provides a molecular basis for their multifaceted role in gene regulation

Our Integrated RBP Regulatory Map reveals numerous cases of combinatorial interactions and multi-functional roles for RBPs, a number of which have been previously described. Such combinatorial interactions appear as off-diagonal groupings in our clustered interaction map (Fig. 2B). For example, FUS, along with several other proteins, is associated with both mRNA splicing and ribosome biogenesis regulatory modules. Both of these functions have been experimentally demonstrated for FUS (Fiesel and Kahle 2011; Moore 2016; Rogelj et al. 2012). Similarly, FASTKD2, depending on the context, is known to play roles in the regulation of mitochondrial mRNA translation and mitochondrial ribosomal biogenesis, respectively (Antonicka and Shoubridge 2015; Popow et al. 2015). Consistently, FASTKD2 is associated with both of these regulatory modules in our integrated map. These examples further highlight the combinatorial nature of RBP function and our ability to effectively capture and separate these interactions.

To go beyond these known examples of multi-functionality and to gain insights into the previously unknown functions, we implemented a label transfer approach for each RBP by analyzing the functions of its closest neighbors in our Integrated RBP Regulatory Map. We observed that, as expected, the closest neighbors often also point to the known functions of RBPs (Fig. 2C). For example, the strong interactions of XRCC6 (Ku70) with DDX51, UTP18, and WDR43 suggest a role in ribosome biogenesis, which was recently demonstrated for this RBP (Shao et al. 2020). Similarly, interactions between PPIL4, NPM1, and SAFB suggest that PPIL4 also takes part in transcriptional regulation. Consistent with this observation, PPIL4 was recently shown to interact with JMJD6, an annotated histone demethylase with known roles in transcriptional control (Barak et al. 2021). Finally, the interaction between LIN28B and RPS3, METAP2, and DDX6 similarly recapitulates its known role in mRNA translation (Basak et al. 2020).

As mentioned above, the use of complementary data types allows our Integrated RBP Regulatory Map to better capture this multifunctionality of RBPs. As a demonstration, for each RBP of interest, we tallied the annotated functions of its closest neighbors across each of our three modalities (Fig. 2D). For instance, when considering ZNF800, we found that UCHL5 and GRWD1, which are among its closest neighbors in the eCLIP dataset, are known chromatin remodeling proteins. However, the proximity labeling data showed that ribosome biogenesis factors, such as DDX21 and RPS11, are also among the proteins in ZNF800’s neighborhood. Consistently, our Integrated RBP Regulatory Map captured both modalities and revealed that the protein neighborhood of ZNF800 consists of both chromatin remodeling and ribosome biogenesis factors (Fig. 2D, middle panel). This showcases that the use of complementary data types allows for the simultaneous capturing of multiple putative RBP functions. TAF15 is another fitting example; eCLIP data suggests that TAF15 is closely related to transcriptional regulators, such as FUS and EWSR1. At the same time, proximity labeling data point to an interaction between TAF15 and the splicing machinery (through SMNDC1 and GPKOW). Perturb-seq data, on the other hand, captures translational regulators PABPC4 and EIF4G2 among TAF15’s close neighbors. Again, the Integrated RBP Regulatory Map for TAF15 contains all of the above-mentioned modules (Fig. 2D, bottom). Our integration procedure can also capture weak interactions that are consistent between modalities. For example, transcription regulators TAF15 and AKAP8L are poorly represented among DGCR8 closest neighbors in the individual data modalities, however, they are included in top-5 RBP’s partners according to the integrated score (Fig. 2D, top) and DGCR8 involvement in chromatin organization as well as its direct interaction with FET proteins has been experimentally shown (Deng et al. 2019; Shiohama et al. 2007).

### Defining functional RBP neighborhoods using BioID-mediated proximity labeling

Having defined the modules that each RBP falls into, we next sought to assign regulatory functions to each of these modules. The proximity labeling data allowed us to go beyond RBP-RBP interactions (Fig. S3A-C) and study the functions of both individual RBPs and their modules by analyzing the totality of their protein neighborhoods. For each RBP, we ranked its neighbors by their enrichment in the biotinylated fraction. We then performed gene-set enrichment analysis (GSEA) to identify the highest over-represented pathways and protein complexes in each RBP neighborhood (Fig. 3A). This procedure allowed us to systematically estimate the involvement of an RBP in a given pathway across all “RBP-pathway” pairs. Conceptually, these estimates reflect the confidence in each annotation, where higher scores denote higher confidence in the proposed association (Fig. S3D). We have visualized the high-confidence annotations in a heatmap along with the major RNA classes that based on our eCLIP analysis are the targets of each RBP module in Fig. 3B.

**Figure 3.**
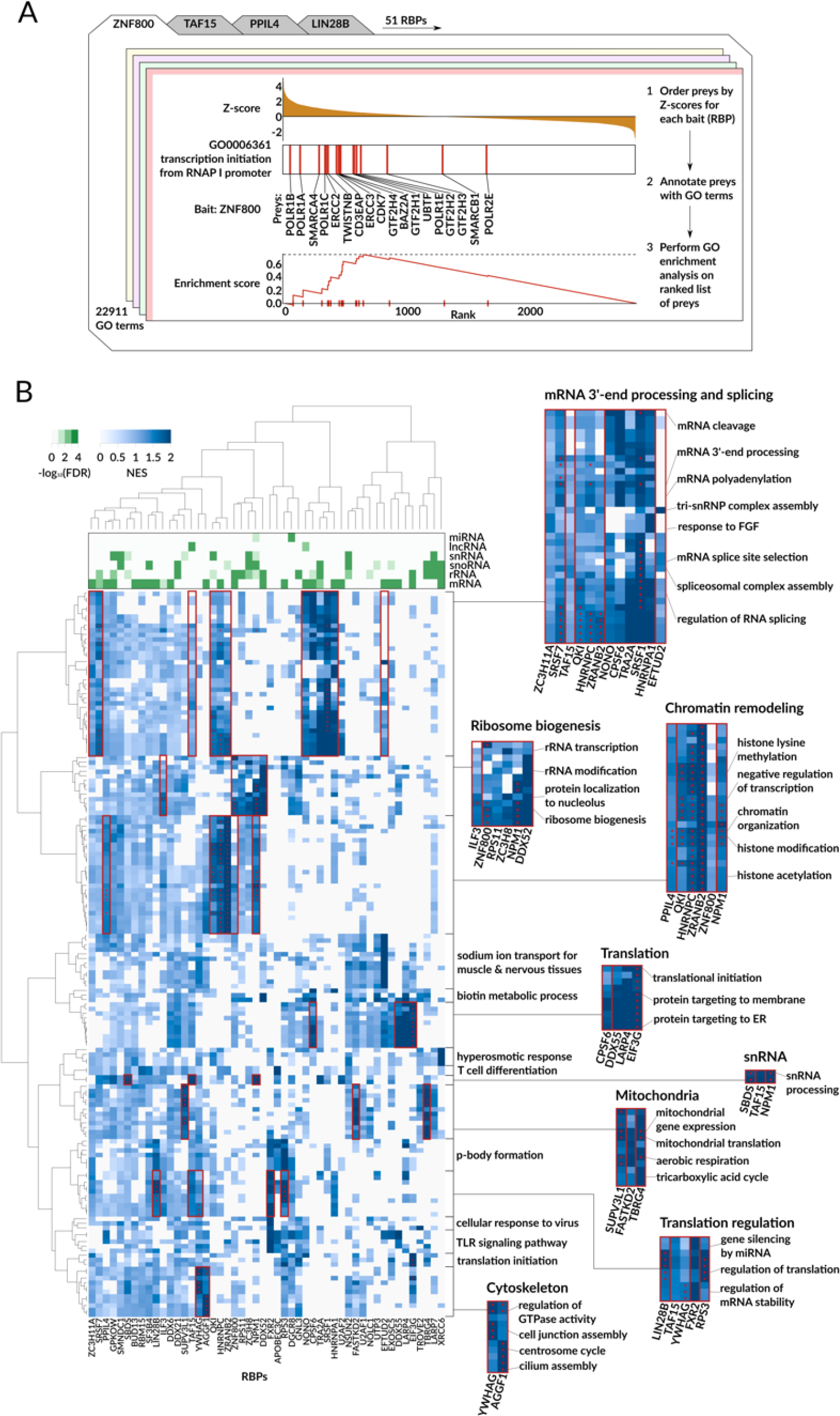
BioID-mediated proximity labeling defines RBP neighborhoods and enables functional annotation of RBPs. **(A)** Overview of our pathway annotation workflow for RBPs. The example provided shows the test for the association of ZNF800 and GO:0006361 (transcription initiation from RNA polymerase I promoter). Proximity-labeled proteins were ranked by their *z*-scores in the ZNF800-BioID dataset, where a higher score implies enrichment relative to control. Experiments were performed in biological triplicates using unlabeled samples as controls (three case vs. three control design). Gene-set enrichment analyses were performed on the resulting ranked list across all RBPs. Each enrichment analysis resulted with p-value and NES score for a given pair of RBP and a pathway. **(B)** A heatmap showing the associations between RBPs and pathways as inferred from proximity labeling data. Columns correspond to the RBPs, rows correspond to individual gene ontology terms (Biological Processes; BP), and the color denotes the GSEA normalized enrichment score (NES). The associations showing FDR < 0.05 are marked with a red asterisk. The green heatmap in the header shows the RBP binding preferences to particular RNA types, as determined based on eCLIP RNA targets. Some known functions of RBPs are highlighted by red boxes and zoomed-in on the right.

In many cases, the previously known functions of RBPs are clearly captured by this approach (Fig. 3B). For example, we have correctly annotated SRSF7, NONO, and HNRNPA1 as splicing-related RBPs that bind predominantly pre-mRNAs. Similarly, we identified RPS11, NPM1, and DDX52 as RBPs that are involved in ribosome biogenesis and directly interact with rRNAs. Our BioID-based annotations also identified RBPs that regulate transcription (HNRNPC, NPM1, QKI) (Mallory et al. 2020; Box et al. 2016; Ren et al. 2021), initiate and regulate mRNA translation (LARP4, EIF3G, RPS3, LIN28B) (R. Yang et al. 2011; des Georges et al. 2015; Basak et al. 2020), participate in snRNA processing (TAF15, NPM1) (Kugel and Goodrich 2009; Nachmani et al. 2019) and mitochondrial metabolism (SUPV3L1, FASTKD2, TBRG4) (A. R. Wolf and Mootha 2014), and modulate centrosome amplification (YWHAG) (Mukhopadhyay et al. 2016).

Our findings also reveal novel and previously unexplored “non-canonical” functions for human RBPs, highlighting the gaps in our current knowledge of RBP annotations that can be systematically addressed with our approach. For example, SRSF7 is primarily known as a splicing factor; however, we observed an equally strong enrichment of mRNA 3’-end processing and polyadenylation pathways, which are not yet annotated in GO but are alluded to in recent publications (Müller-McNicoll et al. 2016; Schwich et al. 2021). Overall, we have annotated 19 RBPs with 1,111 BP GO terms at 5% FDR, of which, 736 (66%) are novel. In the following sections, we have experimentally verified a number of these annotations.

### ZC3H11A and TAF15 are multifunctional post-transcriptional regulators involved in splicing, translation, and RNA stability

We have annotated ZC3H11A and TAF15 as multifunctional RBPs involved in multiple post-transcriptional processes for distinct RNA regulons. ZC3H11A is known to be a part of the TREX complex responsible for mRNA export and was shown to co-localize with SRSF2 in nuclear splicing speckles (Folco et al. 2012; Younis et al. 2018). TAF15, on the other hand, is predominantly studied as part of the TFIID and RNAPII complexes (Jobert, Argentini, and Tora 2009). TAF15’s role as an RNA-binding protein in post-transcriptional control, however, is not as well characterized. Regardless, several studies have shown TAF15 to be involved in stability and processing of lncRNA LINC00665, FGFR4, and GRIN1 mRNAs, and a small subset of other mRNAs in neurons (Ruan et al. 2020; DeJong et al. 2021; Ibrahim et al. 2013; Kapeli et al. 2016). Additionally, it has been shown that TAF15 participates in miRNA-mediated regulation of cell cycle gene expression, and its role in mRNA transport and translation has been suggested based on its pervasive binding to 3’UTR (Ballarino et al. 2013; Kapeli et al. 2016).

Our integrated RBP interaction map suggests that both ZC3H11A and TAF15 are involved in a much wider set of post-transcriptional regulatory processes (Fig. 2). In particular, TAF15 has the highest interaction scores with FUS, SAFB2, EIF4G2, NONO, and SAFB, which in addition to transcription are also associated with translation (EIF4G2) and splicing (NONO). On the other hand, ZC3H11A’s top interacting partners include GPKOW and DHX30, suggesting putative splicing-related functions. Consistently, gene-set enrichment analysis of the proximity-labeling data revealed mRNA export (for ZC3H11A) and transcription (for TAF15), as the highest-scoring pathways. However, we also noted multiple additional high-scoring pathways, including “spliceosomal snRNP assembly” for both RBPs (GO:0000387, NES = 1.5) as well as “mRNA stabilization” (GO:0048255, NES = 1.5) and “positive regulation of translation” (GO:0045727, NES = 1.7) for TAF15 (Fig. 4A).

**Figure 4.**
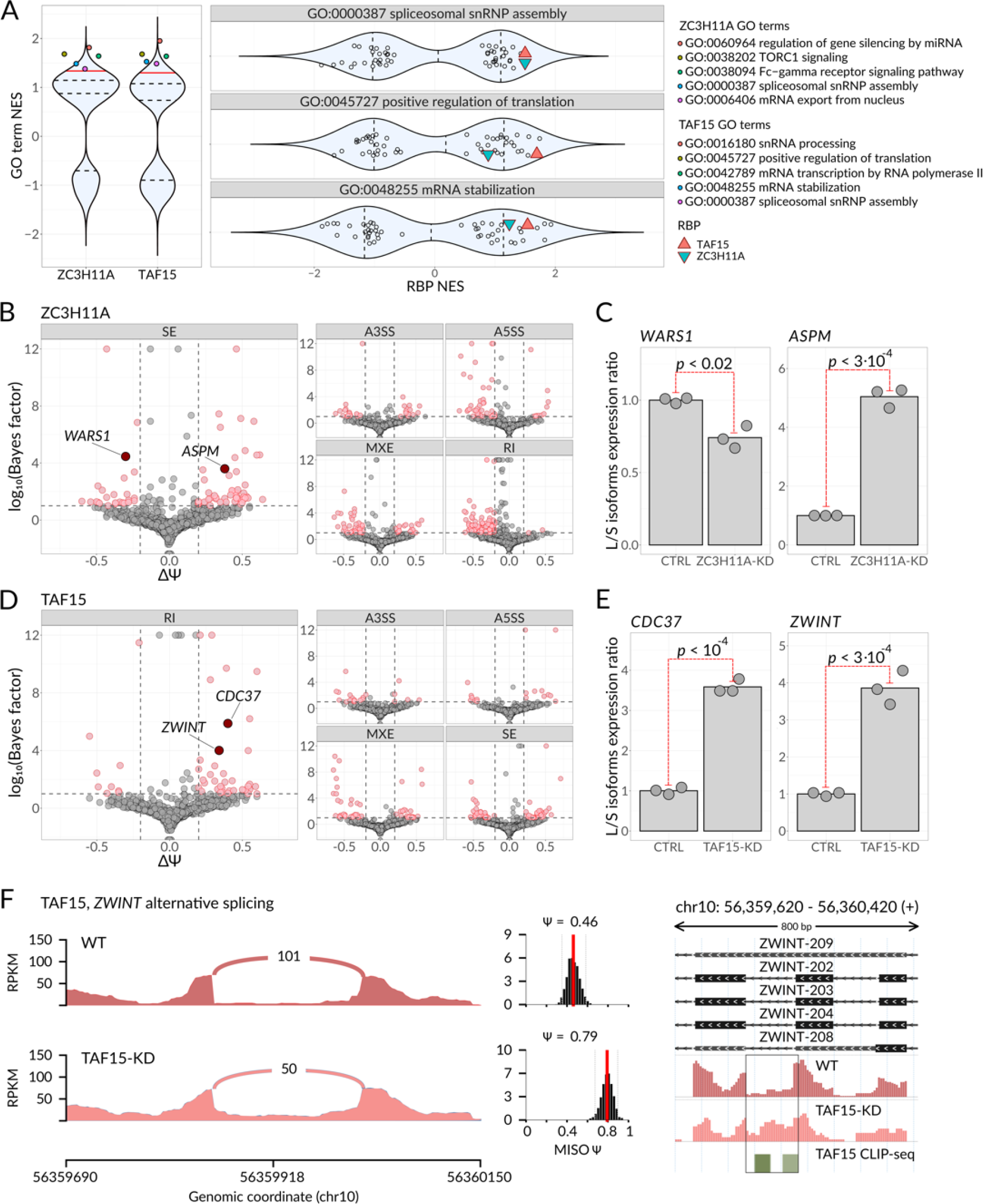
ZC3H11A and TAF15 control multiple independent regulons through distinct regulatory programs. **(A)** Violin plots showing the normalized enrichment scores (NES) resulting from gene set enrichment analysis of proximity labeling data. Left subpanel: NES scores across all the GO-BP terms for ZC3H11A and TAF15 proteins. The five highest scoring pathways are highlighted with color. Right subpanel: NES scores across all studied RBPs for the pathways GO:0000387, GO:0045727, and GO:0048255. ZC3H11A and TAF15 are highlighted with colored triangles. Dashed lines: quartiles; solid red line: 0.9 quantile. **(B)** Scatterplot showing changes in alternative splicing events (ASE) usage upon ZC3H11A knockdown as estimated by MISO. Individual subplots cover different classes of alternative splicing events: Skipped Exon (SE), Retained Intron (RI), Alternative 3’ Splice Site (A3SS), Alternative 5’ Splice Site (A5SS), and Mutually Exclusive Exon (MXE). Dashed lines indicate the following filters: Bayes factor >=10 and the absolute value of isoforms levels difference >=0.2. The ASEs passing these filters are shown in red. **(C)** Relative levels of two skipped exons from the transcripts *WARS1* (left) and *ASPM* (right) were measured by RT-qPCR in control K562 and ZC3H11A-KD cells; n = 3 biological replicates. *P* from t-test performed on log-transformed isoform expression ratios. **(D)** Scatterplot showing changes in alternative splicing events in TAF15 knockdown cells, as in (B). **(E)** Relative levels of two retained introns from the transcripts *CDC37* (left) and *ZWINT* (right) were measured by RT-qPCR in control K562 and ZC3H11A-KD cells; n = 3 biological replicates. *P* from t-test performed on log-transformed isoform expression ratios. **(F)** Left: Sashimi plot illustrating the changes in intron retention event usage in *ZWINT* transcript upon TAF15 knockdown. Right: Genomic view of the *ZWINT* retained intron, RNA-seq profiles from WT and TAF15-KD cells and TAF15 CLIP-seq peaks are shown at the bottom. Y axis: counts per million (CPM). The region corresponding to the alternative splicing event is framed.

To verify these putative roles for ZC3H11A and TAF15 in splicing regulation, we used CRISPR-interference to knockdown these RBPs in K562 cells and performed paired-end total RNA-seq in respective knockdown cells (96% and 98% knockdown efficiency when compared to non-targeting guide RNA, respectively, Fig. S4A-E). Upon silencing either of these genes, we observed a large number of significant alternative splicing events (ASEs) (296 and 190 differentially spliced events for ZC3H11A and TAF15 knockdowns, respectively; Fig. 4B,D). We validated several of these significant ASEs (Fig. S4F-H) using quantitative RT-PCR (Fig. 4C,E); therefore confirming the involvement of ZC3H11A and TAF15 in the regulation of alternative splicing. To confirm that these modulations are the result of direct interaction between ZC3H11A or TAF15 and these target pre-mRNAs *in vivo*, we performed crosslinking and immunoprecipitation followed by sequencing (CLIP-seq) (Licatalosi et al. 2008) for both ZC3H11A and TAF15 in K562 cells. As expected, we observed multiple binding sites in close proximity to the significant ASEs for both ZC3H11A and TAF15 (Fig. 4F, Fig. S4I). Taken together, these results establish ZC3H11A and TAF15 as regulators of alternative splicing for their respective regulons.

In addition to RNA processing and splicing, we also observed a significant and independent association between TAF15 and translational control machinery. To confirm this observation, we performed ribosome footprinting (Ribo-seq; (Ingolia et al. 2012)) as well as matched RNA sequencing in control and TAF15 knockdown cells (Fig. S5A-C). Ribo-seq, which captures active translation, allows us to measure changes in translational efficiency, defined as the ratio of ribosome-protected fragments to total RNA. Consistent with a direct role for TAF15 in translational control, TAF15-bound RNA targets (Van Nostrand et al. 2016) were strongly enriched among mRNAs that were translationally repressed in TAF15 knockdown cells (Fig. 5A). To confirm this observation, we generated and compared protein abundance data in TAF15 KD and control cell lines using quantitative mass spectrometry. As expected, TAF15 targets showed a significant change in their protein abundance without a concomitant change in their mRNA levels (Fig. 5A,B). Taken together, our findings demonstrate that TAF15 moonlights as an enhancer of mRNA translation for its target regulon.

**Figure 5.**
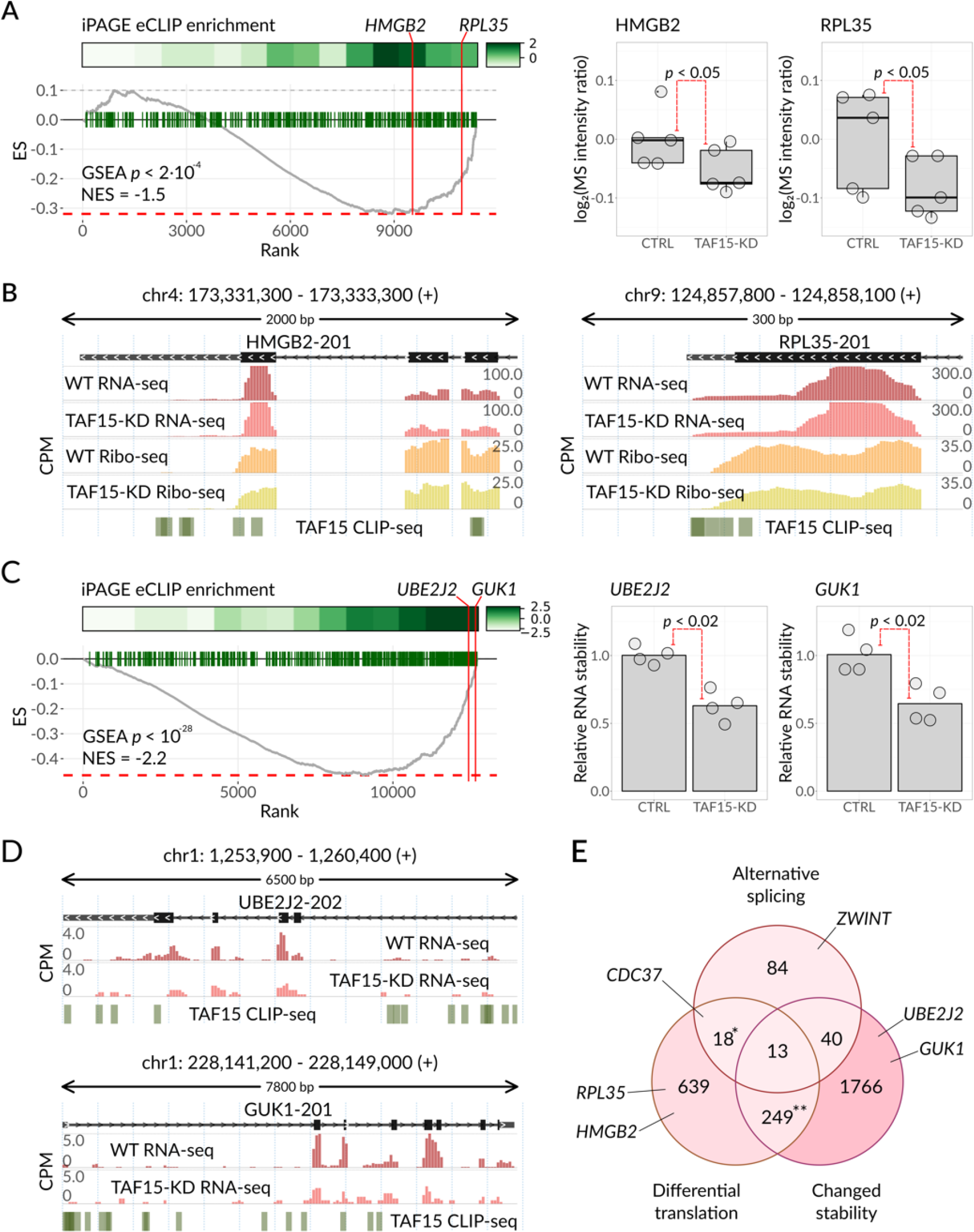
TAF15 is directly involved in RNA translation and stability regulation. **(A)** Left: enrichment analysis of TAF15 mRNA targets among the differentially translated genes (in TAF15-KD cell line compared to WT cell line). The differential ribosome occupancy (RO) measurements in TAF15-KD cells were estimated from Ribo-seq. The genes were sorted based on the RO change (along the x-axis), and the enrichment of TAF15 mRNA targets, inferred from eCLIP data, was calculated using iPAGE (top subpanel) and with GSEA (bottom subpanel, ES stands for the enrichment score). Two example targets, HMGB2 and RPL35, are highlighted. Right: levels of HMGB2 and RPL35 were measured by mass spectrometry in control K562 and TAF15-KD cells. N = 5 biological replicates. *P* from one-sided Wilcoxon rank sum test. **(B)** Genomic view of *HMGB2* (left) and *RPL35* (right). RNA-seq and Ribo-seq WT and TAF15-KD profiles as well as TAF15 CLIP-seq peaks are shown below. Y axis: counts per million (CPM). **(C)** Left: enrichment analysis of TAF15 mRNA targets among the differentially stabilized transcripts (in TAF15-KD cell line compared to WT cell line) measured by α-amanitin treatment. The transcripts were sorted based on stability change (log2FCs). The enrichment of TAF15 RNA targets, inferred from eCLIP data, was calculated with iPAGE (top and middle subpanel) and with GSEA (bottom subpanel). Two example targets, UBE2J2 and GUK1, are highlighted. Right: relative stability of *UBE2J2* and *GUK1* mRNAs were measured as mRNA to pre-mRNA abundances ratio using qPCR in control K562 and TAF15-KD cells. N = 4 biological replicates. *P* from one-sided Wilcoxon rank sum test. **(D)** Genomic view of *UBE2J2* (top) and *GUK1* (bottom). RNA-seq WT and TAF15-KD profiles as well as TAF15 CLIP-seq peaks are shown below. Y axis: counts per million (CPM). **(E)** Venn diagram of TAF15 RNA regulons. Shown are the numbers of genes that exhibit significant changes in splicing (155 genes with Bayes factor >=10), translation (919 genes with FDR < 0.05), or stability (2,068 genes with FDR < 0.05) upon TAF15 knockdown, as captured by RNA-seq, Ribo-seq, and RNA-seq with α-amanitin, respectively. Asterisks reflect the significance obtained from pairwise one-sided Fisher’s exact tests (odds ratio = 2 and *P* < 10^-15^ for translation and stability, odds ratio = 1.5 and *P* < 0.05 for translation and splicing).

We also observed a strong association between TAF15 with regulators of RNA stability including LARP1, SYNCRIP, and RBM10. To further explore this association, we inhibited RNA Polymerase II-mediated transcription with α-amanitin and performed RNA-seq in control and TAF15 knockdown cells in order to estimate changes in mRNA decay upon TAF15 knockdown (Fig. S5D-F) (Lugowski, Nicholson, and Rissland 2018). We then analyzed the changes in mRNA decay among the TAF15 target mRNAs with iPAGE (Goodarzi, Elemento, and Tavazoie 2009) and GSEA (Korotkevich et al. 2021). Both tools demonstrated the enrichment of TAF15-bound RNA targets among those with a shorter half-life in TAF15 knockdown cells (Fig. 5C). To independently verify this observation, we used RT-qPCR to compare mRNA stability of several TAF15 mRNA targets, such as UBE2J2 and GUK1, in TAF15-KD versus control cells (Fig. 5C,D). For all tested targets, we observed significantly lower mRNA stability upon TAF15 knockdown, thus supporting our hypothesis of TAF15 involvement in mRNA stability control.

Our goal in this section was to showcase how a single RBP, in this case, TAF15, can play multiple independent regulatory functions based on the context of its interactions with each regulon. To further explore this notion, we asked whether the three sets of TAF15 target RNAs, corresponding to TAF15 role in splicing, translation, and stability, are in fact distinct and form independent regulons (Fig. 5E). We observed that the translation and stability regulons partially but significantly overlap; this is concordant with the known interdependence of these two biological processes (Radhakrishnan and Green 2016). On the other hand, translation versus splicing and splicing versus stability comparisons show only a small number of targets at their intersections (18 out of 741 and 40 out of 1890 genes, respectively; Fig. 5E). Overall, in K562 cells, TAF15 controls splicing of 155 RNAs, translation of 919 RNAs, and stability of 2068 RNAs; 320 of these RNAs fall into two regulons and only 13 are present in all three pathways. Therefore, the majority of TAF15 mRNA targets are exclusive to each regulon. This further highlights the notion that TAF15 participates in three independent regulatory pathways, each with its own distinct target regulon.

### RNA-binding proteins QKI and ZNF800 are involved in the regulation of transcription

While RNA-binding proteins are often thought to strictly regulate post-transcriptional processes, our integrated RBP interaction map revealed several RNA-binding proteins that are also strongly associated with transcriptional control. Chief among these, we noted ZNF800 and QKI. ZNF800 is a zinc finger protein whose molecular functions are poorly studied, yet it is implicated in diseases such as lung cancer (Zhuo et al. 2020). In contrast, QKI is a well-studied RBP involved in many RNA-related processes and is known to play a major part in neuronal gene regulation and neuron myelination (Chen, Liu, et al. 2021; Chen, Yin, et al. 2021; Zhou et al. 2021; Shin et al. 2021; Åberg et al. 2006).

To assess the potential impact of modulations in ZNF800 and QKI levels on transcriptional activity, we used ATAC-seq to measure changes in chromatin accessibility at or near their DNA binding sites. Based on our proximity labeling results, ZNF800’s protein neighborhood functions in DNA methylation, transcription by RNA polymerase I, rRNA processing, and chromatin remodeling (Fig. 6A). On the other hand, QKI’s neighborhood is associated with histone methylation, RNA splicing, transcription by RNA polymerase II and chromatin organization. To validate the previously unknown role for ZNF800 in chromatin remodeling and confirm recently discovered chromatin-associated QKI functions (Ren et al. 2021), we performed ATAC-seq on control K562 and CRISPRi-generated knockdown cells (Fig. S6A-D) (87% and 76% knockdown, respectively) (Buenrostro et al. 2015). We observed a significant and widespread increase in chromatin accessibility across thousands of regions when these RBPs were silenced (2660 out of 2724 significantly differential regions were upregulated for QKI knockdown, and 1399 out of 1417 significantly differential regions were upregulated in ZNF800 knockdown; Fig, 6B). Among the differentially accessible peaks, the majority are located in close proximity (<1 Kb) of gene promoter regions (Fig. 6B).

**Figure 6.**
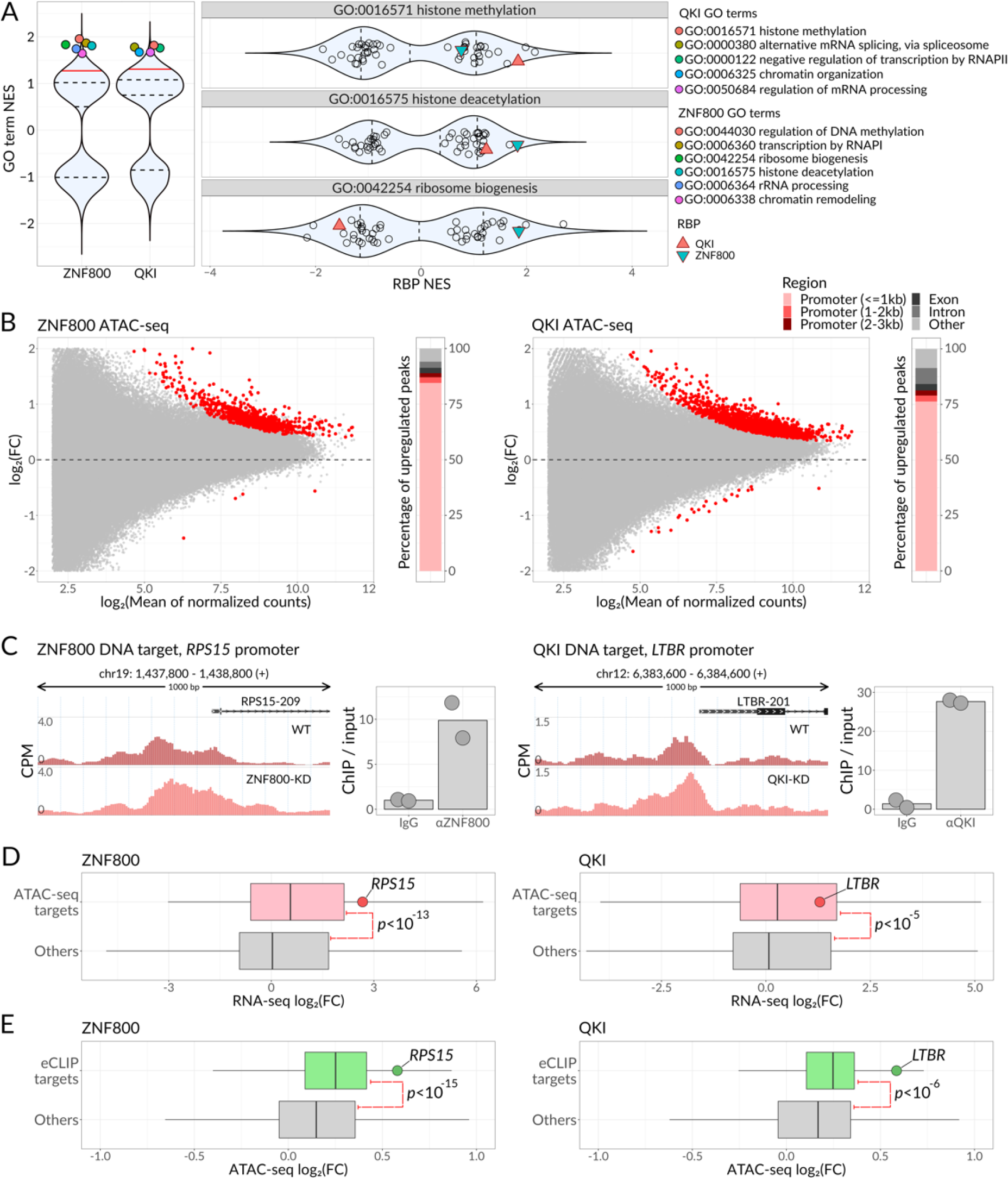
ZNF800 and QKI control gene expression at transcriptional and post-transcriptional levels independently. **(A)** Violin plots showing the normalized enrichment scores (NES) resulting from gene set enrichment analysis of proximity labeling data. Left subpanel: NES scores across all the GO-BP terms for ZNF800 and QKI proteins. The 5 highest scoring pathways are highlighted with color. Right subpanel: NES scores across all the studied RBPs for GO:0016571, GO:0016575 and GO:0042254 GO terms. ZNF800 and QKI are highlighted with colored triangles. Dashed lines: quartiles; solid red line: 0.9 quantile. **(B)** Volcano plots showing differential chromatin accessibility between WT K562 cells and ZNF800-KD (left) or QKI-KD (right) cells. Each point denotes a single ATAC-seq peak; peaks passing 0.1 FDR are colored red. The distribution of peaks among various genomic regions is shown on the right of each volcano plot. **(C)** Genomic view of *RPS15* (left) and *LTBR* (right) promoter regions. ATAC-seq profiles of WT cells along with ZNF800-KD (left) or QKI-KD (right) are shown. Binding of ZNF800 to the *RPS15* promoter region and binding of QKI to the *LTBR* promoter region were measured by ChIP-qPCR in K562 cells and are illustrated on the right of each profile plot. **(D)** Box plots showing the distributions of expression fold changes in WT cells compared to either ZNF800-KD cells (left) or QKI-KD cells (right), as measured by RNA-seq. The distributions for the genes showing significant promoter accessibility changes upon the respective knockdown and for the rest of the genes are shown separately. *P* calculated by one-sided Wilcoxon rank sum test. **(E)** Box plots showing the distributions of chromatin accessibility fold changes in WT cells compared to either ZNF800-KD cells (left) or QKI-KD cells (right), as measured by ATAC-seq. The distributions for ZNF800- or QKI-RNA targets (as defined by eCLIP) and the rest of the genes are shown separately. The top most highly accessible ATAC-seq peak was considered for each gene. P calculated by one-sided Wilcoxon rank sum test.

To further demonstrate that ZNF800 and QKI are chromatin-associated RBPs, we performed ChIP-qPCR on several of their gene targets. Namely, we tested the binding of ZNF800 to the promoter sequences of *RPS15* and *RPL10A*, and the binding of QKI to the promoters of *PRC1* and *LTBR*. As expected, these target sequences were significantly enriched in ChIP samples compared to controls, which demonstrates the localization of ZNF800 and QKI to promoter regions of these identified target genes (Fig. 6C; Fig. S6E,F). In addition to ChIP-qPCR validation, we have tested an overall agreement between the differential ATAC-seq peaks changing upon RBP knockdowns and the published ChIP-exo data (Lai et al. 2021). As expected, ZNF800 and QKI ChIP-exo signal was significantly enriched in differential ATAC-seq peaks compared to the rest of the peaks (U test P < 10^-16^ for ZNF800, Fisher’s exact test odds ratio = 40, P < 10^-16^ for QKI, see Methods).

To test whether the observed changes in chromatin accessibility lead to changes in mRNA expression, we performed RNA-seq in control and QKI- or ZNF800-knockdown cells (Fig. S6A,C). As expected, we observed significantly elevated expression of the genes with increased chromatin accessibility in the ATAC-seq data (Fig. 6D). Together, these observations point to the role of ZNF800 and QKI as transcriptional repressors.

Next, we sought to explore whether the role that ZNF800 and QKI play in transcription inhibition depends on their direct binding to RNA. We tested whether the RNA binding targets of ZNF800 and QKI (based on eCLIP data) correspond to the ATAC-seq peaks that become upregulated upon RBP knockdown. We observed that the genes encoding the RNA binding targets of ZNF800 and QKI contained regions that were significantly more accessible upon the RBP knockdown compared to the other genes (Fig. 6E). This data supports the hypothesis that ZNF800 and QKI achieve their regulatory functions through direct co-transcriptional binding of chromatin-associated RNA.

We have discovered a direct and previously unknown role for QKI and ZNF800 in transcriptional control, even though they were previously thought to be involved in post-transcriptional regulation rather than transcriptional control. Our data suggest that RBP-RNA interactions can often influence transcriptional activity. This further highlights the multimodality and multifunctionality of RNA-binding proteins and the use of the Integrated RBP Regulatory Map for unraveling complex regulatory grammars.

## DISCUSSION

The traditional model of transcriptional control, called the “transcriptional regulatory code” (Harbison et al. 2004) involves *cis*-acting elements such as enhancers and transcription factor binding sites (TFBSs) and *trans*-acting transcription factors (TFs) that bind to these elements in a combinatorial and coordinated manner to create complex regulatory circuits. However, there is no equivalent conceptual framework for studying the combinatorial post-transcriptional control of gene expression. Given that a few hundred RBPs control all aspects of the RNA life cycle, from processing and export to translation and decay, the “one RBP-one function” paradigm does not provide enough complexity to cover all the post-transcriptional regulatory processes that occur in a cell. It is not surprising, then, that RBPs are highly multifunctional and also exhibit a complex and context-specific RNA binding grammar.

Many research initiatives aim to map RBP-bound transcripts as units of post-transcriptional gene expression control. The ENCODE consortium, for example, has conducted a large-scale effort to map protein-DNA and protein-RNA interactions using ChIP-seq and eCLIP, respectively (Van Nostrand et al. 2020). Other groups have also generated similar data modalities, providing a wealth of resources for generating biological networks focused on nucleic acid interactions. However, not all interactions are functional, and not all functions can be generalized across all interactions. This is particularly true for RNA-binding proteins, which often bind thousands of RNAs and have multifaceted and context-dependent regulatory functions across these targets (Fish et al. 2019, 2021). Therefore, it is an oversimplification to consider an RBP regulon as the unit of post-transcriptional control.

Instead of viewing post-transcriptional regulation through the lens of individual RBPs and their bound target RNAs, we propose that the field should instead adopt a more precise definition of RBP regulons that accounts for their context-specificity. To address this issue, we propose the concept of “regulatory modules” as the foundational units of post-transcriptional control, i.e., collections of RNA-binding proteins that work together for a specific function and a distinct target regulon. This approach allows us to capture the many-to-many relationship between RNA-binding proteins and their regulatory functions.

To properly characterize regulatory modules, we must consider both direct and epistatic genetic interactions. The combinatorial action of RBPs may very well be spatially or temporally separate. Therefore, here, we used proximity labeling techniques (BioID labeling and eCLIP) to identify direct physical interactions between RBPs or between RBPs and RNAs, and transcriptome-wide gene expression measurements (Perturb-seq) to identify genetic interactions. Perturb-seq (Adamson et al. 2016) combines the CRISPR-interference technology (Gilbert et al. 2014) to target RBPs and single-cell RNA sequencing to capture changes in the transcriptomic state of the cell in response to RBP knockdowns (Fig. S3E-G). This rich data allowed us to model the genetic interactions manifold, which describes the transcriptional states a cell can occupy upon perturbation (Norman et al. 2019).

To capture the protein neighborhood of each RBP, we used BioID labeling-based pulldown followed by mass spectrometry (D. I. Kim et al. 2016). Proximity-based covalent biotin labeling techniques improve upon classic antibody-based pulldown methods by relying on the strong biotin-streptavidin interaction, which allows for harsh washes and therefore tends to have low levels of nonspecific binding (P. Li et al. 2017) (Roux et al. 2018). Biotin labeling techniques are well-suited for surveying the spatial proximity between proteins within a narrow diameter (Gingras, Abe, and Raught 2019). To date, only a few large-scale datasets of dozens or hundreds of bait proteins have been generated (Antonicka et al. 2020; Go et al. 2021; Youn et al. 2018). In this study, we extended these efforts by applying proximity-dependent biotinylation (BioID) analysis to 50 human RBPs with a well-controlled and rigorous design. We validated the expression of individual fusion proteins by Western blotting (Fig. S3A,B) and found that including matched pulldown controls for every fusion protein significantly improved the quality of our data, as pulldown mass spectrometry profiles were generally more similar to their matched negative controls than to other pulldown profiles (Fig. S3C).

In sum, our Integrated RBP Regulatory Map reveals that multi-functional RBPs playing different and even divergent roles depending on their specific context are the rule, rather than the exception. Therefore, studying the role of RBPs in gene expression regulation requires a substantially improved understanding of higher-order combinatorial interactions between these proteins. We believe that this study takes a step towards building a systematic and integrative framework for tackling this problem.

## DATA AND CODE AVAILABILITY

All the sequencing data is available at Gene Expression Omnibus (GEO), identifier GSE225809. All of the original code is available at https://github.com/goodarzilab/RBP_modules.

## ACKNOWLEDGEMENTS

The authors thank Artemii Bakulin and Heather Karner for helpful discussions. A.N. was supported by DoD PRCRP Horizon Award W81XWH-19-1-0594. D.M. was supported by an MD fellowship from the Boehringer Ingelheim Fonds.

## AUTHOR CONTRIBUTIONS

M.K., F.M., and H.G. designed the study. M.K. and J.Y. performed PerturbSeq experiments.

M.K, A.N, F.T., A.D., R.B., M.D., F.M. performed proximity labeling experiments. M.K, F.T., A.D. and D.M. performed western blotting experiments. A.B. and I.K. performed a re-analysis of ENCODE eCLIP data. M.K., A.B. and I.K. performed the dataset integration. M.K., A.B. and I.K. performed the RBP functional annotation. M.K. performed CRISPRi knockdown experiments. M.K. and B.C. performed RNA-seq experiments. M.K., H.M. and R.C. performed ATAC-seq experiments. K.G. and H.G. performed ChIP-qPCR experiments. D.M. and H.G. performed qPCR experiments. T.J. and B.C. performed α-amanitin treatment experiments. A.N. performed ribosome profiling experiments. M.K., A.B., S.M., V.S. and H.G. performed data analysis. S.Z. performed CLIP-seq experiments. M.K., A.B., I.K. and H.G. wrote the manuscript with input from all authors.

## STAR METHODS

### Cell culture

All cells were cultured in a 37°C 5% CO2 humidified incubator. The 293T cells (ATCC

CRL-3216) were cultured in DMEM high-glucose medium supplemented with 10% FBS, glucose (4.5 g/L), L-glutamine (4 mM), sodium pyruvate (1 mM), penicillin (100 units/mL), streptomycin (100 μg/mL) and amphotericin B (1 μg/mL) (Gibco). The K562 cell line was cultured in RPMI-1640 medium supplemented with 10% FBS, glucose (2 g/L), L-glutamine (2 mM), 25 mM HEPES, penicillin (100 units/mL), streptomycin (100 μg/mL) and amphotericin B (1 μg/mL) (Gibco). All cell lines were routinely screened for mycoplasma with a PCR-based assay.

### Western blotting

Cell lysates were prepared by lysing cells in ice-cold RIPA buffer (25 mM Tris-HCl pH 7.6, 0.15 M NaCl, 1% IGEPAL CA-630, 1% sodium deoxycholate, 0.1% SDS) containing 1X protease inhibitors (Thermo Scientific). Lysate was cleared by centrifugation at 20,000 x g for 10 min at 4°C. Samples were denatured for 10 min at 70°C in 1X LDS loading buffer (Invitrogen) and 50 mM DTT. Proteins were separated by SDS-PAGE using 4-12% Bis-Tris NuPAGE gels, transferred to nitrocellulose (Millipore), blocked using 5% BSA, and probed using target-specific antibodies. Bound antibodies were detected using dye-conjugated secondary antibodies (Licor) according to the manufacturer’s instructions. Antibodies: HA (BioLegend 901533), eIF3I (BioLegend 646701), beta-tubulin (Proteintech 10094-1-AP), GAPDH (Proteintech 10494-1-AP).

### RNA isolation

Total RNA for RNA-seq and RT-qPCR was isolated using the Zymo QuickRNA isolation kit with in-column DNase treatment per the manufacturer’s protocol.

### RNA treatment with α-amanitin

K562 and K562 TAF15 knockdown cell lines were seeded at 1M/1mL density in 2 replicates. Cells were infected with 10 μg/mL α-amanitin(Sigma-Aldrich A2263) for 8-9 h prior to total RNA extractions. Total RNA for downstream RNA-seq was isolated using Zymo QuickRNA Microprep isolation kit with in-column DNase treatment per the manufacturer’s protocol.

### RNA-seq

RNA-seq libraries were prepared using SMARTer Stranded Total RNA-Seq Kit v2 - Pico Input Mammalian (Takara).

### CRISPRi-mediated gene knockdown

K562 cells expressing dCas9-KRAB fusion protein were constructed by lentiviral delivery of pMH0006 (Addgene #135448) and FACS isolation of BFP-positive cells.

The lentiviral constructs were co-transfected with pCMV-dR8.91 and pMD2.D plasmids using TransIT-Lenti (Mirus) into 293T cells, following the manufacturer’s protocol. The virus was harvested 48 hours post-transfection and passed through a 0.45 µm filter. Target cells were then transduced overnight with the filtered virus in the presence of 8 µg/mL polybrene (Millipore).

Guide RNA sequences for CRISPRi-mediated gene knockdown were cloned into pCRISPRia-v2 (Addgene #84832) via BstXI-BlpI sites. After transduction with sgRNA lentivirus, K562 cells were selected with 2 µg/mL puromycin (Gibco). Knockdown of target genes was assessed by RT-qPCR using PerfeCTa SYBR Green SuperMix (QuantaBio) per the manufacturer’s instructions. HPRT1 was used as endogenous control.

### BioID2-RBP fusion cell line generation

50 RBPs were selected based on the 2 criteria: (i) ENCODE eCLIP data availability for a given RBP, (ii) presence of a given RBP in the ORFeome entry clone library (X. Yang et al. 2011). In order to construct the cell lines stably expressing BioID2-RBP fusion proteins, we first cloned in an open reading frame of BioID2 enzyme (D. I. Kim et al. 2016), followed by a linker (YPAFLYKVVYGGGGSGGGGSGGGGS) and attR-flanked *ccdB* counterselection marker for Gateway cloning, into the pWPI backbone (Addgene #12254). We then used Gateway LR Clonase II Enzyme mix (Thermo Fisher) to clone the open reading frames of the RBPs of interest (from ORFeome entry clone library (X. Yang et al. 2011)) into the destination vector. The lentiviral constructs were co-transfected with pCMV-dR8.91 and pMD2.D plasmids using NanoFect (ALSTEM) into 293T cells, following manufacturer’s protocol. The virus was harvested 48 hours post-transfection and passed through a 0.45 μm filter. K562 cells were then transduced for 2 hours while centrifuging (800 RPM) with the filtered virus in the presence of 8 μg/mL polybrene (Millipore). Cells were selected with 20 μg/mL blasticidin for 5 days (Gibco). The expression of the fusion protein was validated by western blotting.

### Biotin treatment and pulldown

The pulldown was performed as described in (D. I. Kim et al. 2016). Cells were incubated with biotin-depleted media (biotin-free RPMI-1640 medium, supplemented with 10% dialyzed FBS, glucose (2 g/L), L-glutamine (2 mM), 25 mM HEPES, penicillin (100 units/mL), streptomycin (100 μg/mL) and amphotericin B (1 μg/mL) for 72 h before analysis. For BioID2 pulldown, 12 × 10^6^ cells per replicate were incubated with 50 μM biotin for 24 h. For the negative control samples, 12 × 10^6^ cells per replicate were incubated with DMSO. After three times of PBS wash, the cells were lysed in 1 ml of lysis buffer containing 50 mM Tris, pH 7.5, 150 mM NaCl, 1 mM EDTA, 1 mM EGTA, 1% Triton X-100, 1% Sodium deoxycholate, 0.1% SDS, 1 × Complete protease inhibitor (Halt Phosphatase Inhibitor Cocktail; Life Technologies), and 250 units benzonase (EMD millipore). The lysates were passed through a 25G needle 10 times and cleared 10 min at 14,000 g at +4°C. The protein concentration was measured with BCA Protein Assay Kit (Thermo Scientific); the lysate was diluted to a concentration of 2 μg/mL. 500 μl of lysate was incubated with 125 μl of Dynabeads (MyOne Streptavidin C1; Life Technologies) overnight with rotation at +4°C. Beads were collected using a magnetic stand and washed twice with 2% (wt/vol) SDS, twice with wash buffer containing 50 mM Tris, pH 7.5, 500 mM NaCl, 1 mM EDTA, 1 mM EGTA, 1% Triton X-100, 0.1% SDS, twice with wash buffer containing 50 mM Tris, pH 7.5, 150 mM NaCl, 1 mM EDTA, 1 mM EGTA, 1% Triton X-100, 0.1% SDS, then boiled for 5 min in 50 μl of elution buffer containing 2% SDS, 100mM DTT, Tris-HCl pH 7.5. The supernatant was collected and saved for mass spectrometry analysis.

### Mass spectrometry analysis

Eluted BioID samples were reduced by the addition of 100 mM DTT and boiling at 95°C for 10 min, before being subjected to Filter Aided Sample Preparation (FASP) (Wisniewski et al., Anal. Biochem., 2011) to generate tryptic peptides, as described previously (Dermit et al, Dev Cell, 2020). Briefly, samples were diluted 7-fold in UA buffer (8M urea, 100 mM Tris HCl pH 8.5), transferred to Vivacon 500 Hydrosart centrifugal filters with a molecular cut-off of 30kDa (Sartorius), and concentrated by centrifugation at 14,000 g for 15 min. Filters were then washed twice by addition of 0.2 mL of UA buffer to the filter tops and re-concentrating. Reduced cysteine residues were then alkylated by addition of 100µL of 50 mM iodoacetamide dissolved in UA buffer, and incubation at room temperature in the dark for 30 min. The iodoacetamide solution was then removed by centrifugation at 14,000 g for 10 min, and samples were washed twice with 0.2 mL of UA buffer as before. Urea was then removed from samples by performing three washes with 0.2 mL of ABC buffer (0.04 M ammonium bicarbonate). Filters were then transferred to fresh collection tubes, and proteins were digested by addition of 0.3 µg of MS grade Trypsin (Sigma-Aldrich) dissolved in 50 µL of ABC buffer, and overnight incubation in a thermo-mixer at 37°C with gentle shaking (600 rpm). The resulting peptides were eluted from the filters by centrifugation at 14,000 g for 10 min. Residual remaining peptides were further eluted by addition of 100 µL ABC to the filter tops and centrifugation. This was repeated once and the combined eluates were then dried by vacuum centrifugation (no heating) and reconstitution in 2% Acetonitrile (ACN), 0.2% Trifluoroacetic acid (TFA), followed by desalting using C18 StageTips (Rappsilber, et al., Nat Protoc. 2007). The desalted peptides were dried again by vacuum centrifugation (45°C) and re-suspended in A* buffer (2%ACN, 0.5% Acetic acid, 0.1% TFA in water) before LC-MS/MS analysis. 1/3rd of each sample was analyzed on a Q-Exactive plus Orbitrap mass spectrometer coupled with a nanoflow ultimate 3000 RSL nano HPLC platform (Thermo Fisher). Samples were resolved at a flow rate of 250 nL/min on an Easy-Spray 50 cm x 75 μm RSLC C18 column with 2 µm particle size (Thermo Fisher), using a 123 minutes gradient of 3% to 35% of buffer-B (0.1% formic acid in ACN) against buffer-A (0.1% formic acid in water), and the separated peptides were infused into the mass spectrometer by electrospray (1.95 kV spray voltage, 255°C capillary temperature). The mass spectrometer was operated in data-dependent positive mode, with 1 MS scan followed by 15 MS/MS scans (top 15 method). The scans were acquired in the mass analyser at 375-1500 m/z range, with a resolution of 70,000 for the MS and 17,500 for the MS/MS scans. A 30-second dynamic exclusion of fragmented peaks was applied to limit repeated fragmentation of the same ions.

### Perturb-seq

68 RBPs were chosen for Perturb-seq analysis based on the clustering analysis of ENCODE eCLIP dataset and DeepBind dataset (Alipanahi et al. 2015). Perturb-seq experiment was performed as previously described (Datlinger et al. 2017). Briefly, a library of 205 sgRNAs (5 non-targeting sgRNAs and 200 sgRNAs targeting 100 genes, 2 sgRNAs per gene) was ordered as a pooled oligonucleotide library from Twist Bioscience with the following design:

[ATCTTGTGGAAAGGACGAAACACCG]-[Protospacer Sequence]-[GTTTTAGAGCTAGAAATAGCAAGTTAAAATAAGGC]

The library was PCR-amplified using Q5 Hot Start High-Fidelity 2X Master Mix (NEB) with the primers with the following sequences: 5’-ATCTTGTGGAAAGGAC-3’ and 5’-GCCTTATTTTAACTTGCTA-3’. To clone libraries into CROPseq-Guide-Puro vector (Addgene 86708), the starting vector was digested with BsmBI following the protocol outlined in (Sanjana, Shalem, and Zhang 2014). The library was cloned into the digested backbone using Gibson Assembly method (Thomas, Maynard, and Gill 2015). The reaction product was transformed into Takara Stellar competent cells according to manufacturer recommendations, grown overnight in 100 mL LB with ampicilin and purified using ZymoPURE II Plasmid Midiprep Kit. K562 cells were infected with the plasmid library at a low multiplicity of infection to minimize double infection. The infected cells were selected with 2 µg/mL puromycin for 3 days. Live cells were isolated on a flow cytometer (FACSAria II) by propidium iodide staining. Approximately 5000 live cells were captured by 10X Chromium Controller using Chromium Single Cell 3’ Reagent Kits v2. Sample preparation was performed according to the manufacturer’s protocol. Samples were sequenced on a NovaSeq 6000 using the following configuration: Read 1: 28, i7 index: 8, i5 index: 0, Read 2: 98.

To facilitate sgRNA assignment, sgRNA-containing transcripts were additionally amplified by PCR reactions by modifying a previously published approach (Hill et al. 2018). The following primers were used for amplification: 5’-AATGATACGGCGACCACCGAGATCTACAC-3’ and 5’-CAAGCAGAAGACGGCATACGAGATTACGACAGGTGACTGGAGTTCAGACGTGTGCTCTTCCGATCTggactatcatatgcttaccgtaacttgaaag-3’. PCR product was cleaned up by 1.0x SPRI beads (SPRIselect; BECKMAN COULTER; Cat. No. B23317). Samples were sequenced using paired-end 150 bp sequencing on an Illumina MiSeq sequencer.

### Ribosome profiling

Ribosome profiling was performed as previously described (McGlincy and Ingolia 2017). Briefly, approximately 10x10^6^ cells were lysed in ice-cold polysome buffer (20 mM Tris pH 7.6, 150 mM NaCl, 5 mM MgCl_2_, 1 mM DTT, 100 µg/mL cycloheximide) supplemented with 1% v/v Triton X-100 and 25 U/mL Turbo DNase (Invitrogen). The lysates were triturated through a 27G needle and cleared for 10 min at 21,000 x g at 4°C. The RNA concentration in the lysates was determined with the Qubit RNA HS kit (Thermo). Lysate corresponding to 30 µg RNA was diluted to 200 µl in polysome buffer and digested with 1.5 µl RNaseI (Epicentre) for 45 min at room temperature. The RNaseI was then quenched by 10 µl SUPERaseIN (Thermo).

Monosomes were isolated using MicroSpin S-400 HR (Cytiva) columns, pre-equilibrated with 3 mL polysome buffer per column. 100 µl digested lysate was loaded per column (two columns were used per 200 µl sample) and centrifuged 2 min at 600 x g. The RNA from the flow-through was isolated using the Zymo RNA Clean and Concentrator-25 kit. In parallel, total RNA from undigested lysates was isolated using the same kit.

Ribosome-protected footprints (RPFs) were gel-purified from 15% TBE-Urea gels as 17-35 nt fragments. RPFs were then end-repaired using T4 PNK (NEB) and pre-adenylated barcoded linkers were ligated to the RPFs using T4 Rnl2(tr) K227Q (NEB). Unligated linkers were removed from the reaction by yeast 5’-deadenylase (NEB) and RecJ nuclease (NEB) treatment. RPFs ligated to barcoded linkers were pooled, and rRNA-depletion was performed using riboPOOLs (siTOOLs) as per the manufacturer’s recommendations. Linker-ligated RPFs were reverse transcribed with ProtoScript II RT (NEB) and gel-purified from 15% TBE-Urea gels. cDNA was then circularized with CircLigase II (Epicentre) and used for library PCR. First, a small-scale library PCR was run supplemented with 1X SYBR Green and 1X ROX (Thermo) in a qPCR instrument. Then, a larger scale library PCR was run in a conventional PCR instrument, performing a number of cycles that resulted in ½ maximum signal intensity during qPCR. Library PCR was gel-purified from 8% TBE gels and sequenced on a SE50 run on Illumina HiSeq4000 instrument at UCSF Center for Advanced Technologies.

### ATAC-seq

The assay for transposase-accessible chromatin using sequencing (ATAC-seq) was performed according to the optimized Omni-ATAC protocol (Corces et al. 2017; Grandi et al. 2022). Briefly, samples containing 50,000 cells as input were pelleted, lysed, washed, and re-pelleted using the lysis and wash buffers specified in the Omni-ATAC protocol. A transposition mix containing Tn5 was then added to the samples, and the transposition reaction was carried out for 30 minutes at 37 °C in a thermomixer with 1000 rpm mixing. After transposition, the transposed DNA was purified using the DNA Clean & Concentrator-5 Kit (Zymo Research, cat. no. D4014).The samples underwent two PCR steps. First, a pre-amplification was performed for 3 cycles to attach unique barcoded adapters to the transposed DNA sample. The concentration of each pre-amplified sample was quantified via qPCR using the NEBNext Library Quant Kit (New England Biolabs, cat. no. E7630). Afterward, samples underwent a second PCR amplification step to obtain the desired DNA concentration of 8 nM in 20 ul. DNA cleanup and qPCR quantification were performed again, and final libraries were diluted down to exactly 8 nM using sterile water. Samples were sequenced using paired-end 75-bp sequencing on an Illumina NextSeq sequencer.

### ChIP-qPCR

ChIP-qPCR was performed as described in (Rossi, Lai, and Pugh 2018).

Human chronic myelogenous leukemia K562 cells were grown at 37 °C and 5% CO2 in RPMI-1640 medium supplemented with 10% FBS, glucose (2 g/L), L-glutamine (2 mM), 25 mM HEPES, penicillin (100 units/mL), streptomycin (100 μg/mL) and amphotericin B (1 μg/mL) (Gibco). 20 million cells per sample were washed with PBS (in duplicate), pelleted, and cross-linked with 1% paraformaldehyde for 10 minutes at room temperature. Glycine at a final concentration of 125mM was added to the samples and incubated at room temperature for 5 minutes to quench the paraformaldehyde. Samples were washed with PBS, pelleted, flash-frozen, and stored at -80. Samples were thawed, lysed in 200 µl Membrane Lysis Buffer (10 mM Tris-HCl pH 8.0, 10 mM NaCl, 0.5% IGEPAL CA-630, 1X protease inhibitors), and incubated on ice for 10 minutes. Samples were centrifuged at 4°C at 2500 g for 5 minutes, resuspended in 200 µl Nuclei Lysis Buffer (50 mM Tris pH 8.0, 10 mM EDTA, 0.32% SDS, 1X protease inhibitors), and incubated on ice for 10 minutes. 120 µl of IP Dilution Buffer (20 mM Tris-HCl pH 8.0, 2 mM EDTA, 150 mM NaCl, 1% Triton X-100, 1X protease inhibitors) was added to the samples, and the samples were sonicated using the Bioruptor UCD-200 sonicator for 7 minutes with 30 second on/off intervals for a total of 3 times. Samples were centrifuged at 4°C at 21000 g for 5 minutes to clear the lysate, and the supernatant containing the chromatin was stored at -80.

230 µl IP Dilution Buffer was added to 270 µl chromatin along with 3 µg ZNF800 or QKI antibody or same species IgG, and the samples were incubated overnight at 4°C. The next day, the ChIP samples were spun downat 4°C at 16000 g for 5 minutes and the supernatant was transferred onto 20 µl of washed Protein A/G beads (Pierce). Samples were incubated for 2 hours at 4°C.

The ChIP material was washed once with 500 µl of cold FA lysis low salt buffer (50 mM Hepes-KOH pH 7.5, 150 mM NaCl, 2 mM EDTA, 1% Triton-X 100, 0.1% sodium deoxycholate), twice with cold NaCl high salt buffer (50 mM Hepes-KOH pH 7.5, 500 mM NaCl, 2 mM EDTA, 1% Triton-X 100, 0.1% sodium deoxycholate), once with cold LiCL buffer (100 mM Tris-HCl pH 8.0, 500 mM LiCl, 1% IGEPAL CA-630, 1% sodium deoxycholate), and twice with cold 10 mM Tris 1 mM EDTA pH 8.0. Samples were eluted in 300 µl of Proteinase K reaction mix (20 mM Tris pH 8, 300 mM NaCl, 10 mM EDTA, 5 mM EGTA, 1% SDS, 60 µg Proteinase K) and incubated at 65°C for 1 hour. The supernatant was transferred to phase lock tubes (VWR), purified via phenol-chloroform extraction, and eluted in 30 µl 10 mM Tris pH 8.0.

qPCR was performed using PerfeCTa SYBR Green SuperMix (QuantaBio) per the manufacturer’s instructions. HPRT1 was used as endogenous control.

### Crosslinking and Immunoprecipitation

K562 cells were harvested and crosslinked with ultraviolet radiation (400 mJ/cm2). Cell lysates were then treated with high (1:3000 RNase A and 1:100 RNase I) and low dose (1:15000 RNase A and 1:500 RNase I) of RNase A and RNase I separately and combined after treatment. Antibodies to TAF15 (Thermo MA3-078) or ZC3H11A (Abcam ab241612) was first conjugated to protein A/G beads (Pierce) and then added to cell lysates to immunoprecipitate protein-RNA complex. This was followed by on beads dephosphorylation, polyadenylation and IRDye® 800CW DBCO Infrared Dye (LI-COR) end labeling of the immunoprecipitated RNA fragments. RNA-protein complex was then resolved by SDS-PAGE and visualized on nitrocellulose membrane.

Membranes were then cut and treated with proteinase K to release RNA. We then used Takara smarter small RNA sequencing kit reagents with a custom UMI-oligo dT primer (CAAGCAGAAGACGGCATACGAGATNNNNNNNNGTGACTGGAGTTCAGACGTGTGCTCTTCCGATCTTTTTTTTTTTTTTT) to synthesize cDNA. Sequencing libraries were then prepared with SeqAmp DNA Polymerase (Takara) and sequenced on an illumina Hiseq 4000 sequencer.

### Computational Tools

#### Reanalysis of enhanced CLIP ENCODE data

To reliably identify RNA targets of RBPs in K562 cells, we started with the raw eCLIP FASTQ files of ‘released’ K562 experiments for 120 RBPs that were available in the ENCODE database. The analysis was performed as follows: (1) the reads were preprocessed in the same way as in (Van Nostrand et al. 2016) including adapter trimming with *cutadapt* (v1.18) (Martin 2011), (2) preprocessed reads were mapped to the hg38 genome assembly with GENCODE v31 comprehensive annotation using *hisat2* (v.2.1.0) (D. Kim et al. 2019), (3) the aligned reads were deduplicated using the barcodecollapsepe.py script (https://github.com/YeoLab/eclip/tree/master/bin) as in (Van Nostrand et al. 2016), (4) properly paired and uniquely mapped second reads were extracted using *samtools* (v.1.9, with -f 131 -q 60 parameters) (H. Li et al. 2009), (5) gene-level read counts were obtained with *plastid* (v.0.4.8) by counting 5’ ends of the reads (Dunn and Weissman 2016), (6) analysis of specific enrichment against size-matched control experiments was performed with *edgeR* (v.3.18.1) for each RBP separately, considering only genes passing 2 cpm in at least 2 of 3 samples (Robinson, McCarthy, and Smyth 2010). Reliable RNA targets of each RBP were defined as those passing 5% FDR and log_2_(Fold Change) > 0.5, see Supplementary Table S8. eCLIP target scores (TSs) used in datasets integration were estimated as -log10(P)*sign(logFC) for every “RBP-gene” pair separately.

#### Gene set enrichment analysis of RBP RNA targets

A joint set of 22471 genes detected at 2 cpm in at least two samples of one eCLIP experiment was used as the background for further analysis. RBPs preferences to bind RNAs of a particular type were assessed using a one-sided Fisher’s exact test. The following types of RNAs were selected based upon GENCODE annotation: miRNA, lncRNA, protein_coding, snRNA, snoRNA, rRNA. For each RBP separately, the p-values were adjusted for multiple testing using FDR correction for the number of tested RNA types.

Visualization of the eCLIP, RNA-seq, and ATAC-seq profiles generated using *bedtools genomecov* (v.2.27.1) was performed with *svist4get* (v.1.2.24) (Quinlan and Hall 2010; Egorov et al. 2019).

#### Functional annotation of RBPs

To annotate the RBPs based upon preys identified in BioID experiments, target scores (TSs) were estimated as -log10(P)*sign(log-FoldChange) for every bait-prey pair separately. Next, for each prey, TSs were converted to Z-scores by estimating mean and average across baits. The preys were ranked by Z-scores and Fgsea R package (v.1.12.0) was applied to perform gene set enrichment analysis with 100000 permutations and three GO terms annotation sets (BP, MF, and CC, each taken separately) (Korotkevich et al. 2021). The annotation sets were generated with the go.gsets function of *gage* R package (v.2.36.0) (Luo et al. 2009). Lists of 2865 Entrez ids of preys were used in *fgsea* analysis for each RBP of the total set of fifty. GO terms with NES > 2 for at least one RBP were considered when plotting Figure 3 and Figure S3 (related GO terms were merged manually), negative NES were zeroed for clarity and easier interpretability of the consequent clusterization, see complete data in Table S3). Ward.D2 clusterization along with cosine distance (1 - cosine similarity) were used to generate the heatmaps using the heatmap.2 function of the *gplots* R package (v.3.1.1) (Warnes et al. 2009).

To check the consistency between predicted and known RBP annotations, the same procedure was performed excluding the Z-scoring step to avoid penalizing common generic GO terms e.g. “organelle”, “cell”, etc. The resulting GSEA p-values and NESs were used to calculate the <RBP, GO term> scores as -log10(P)*sign(log-FoldChange) for each RBP and GO term separately. RBPs’ “true” annotations were extracted from the same GO BP, CC, or MF annotation set as used in GSEA. Finally, all data were merged to generate the ROC curve with *PRROC* (v.1.3.1) roc.curve function (Grau, Grosse, and Keilwagen 2015).

#### Dataset integration

The functional similarity of RBPs was estimated by joint analysis of eCLIP, BioID, and Perturb-seq data (Figure 1, Figure S1). First, TS Z-scores were calculated for every gene across RBPs separately for each type of experimental data (eCLIP, BioID, or Perturb-seq) in the same way as preys of the BioID data, see above and Figure S1(1). Next, cosine distance was computed for all 7140 pairs of different RBPs followed by ranking and calculation of empirical p-values defined as a fraction of RBP pairs with the cosine distance less than the score of the tested pair, see Figure S1(2). The empirical p-values were aggregated with logitp function from the *metap* R package (v.1.4), see Figure S1(3) and S1(4), raw (non-aggregated) p-values were used for the RBP pairs assessed in a single type of experiment (George and Mudholkar 1983). Heatmap.2 function of the *gplots* R package (v.3.1.1) with cosine distance and Ward’s (ward.D2) clusterization was used to generate the integration heatmap shown in Figure 2.

STRING-based RBP interaction heatmap was generated using protein links’ combined scores (STRING v.11.5) and the same RBP clusterization received from the integration procedure (Szklarczyk et al. 2018). To test the overlap between STRING and integrated RBP interactions, we performed one-sided Fisher’s exact test on 2x2 contingency tables built by thresholding integrated distances below certain quantile and comparing to those with STRING score over 700 (high confidence interactions according to STRING, Figure S2E). Only 3403 RBP pairs with the estimated STRING score, ranging from 150 to 999, among 7140 total pairs were used as the superset. To estimate and compare RBP module sizes that can be assembled using the integrated distances or the STRING annotation, RBP-centered modules were obtained for different distance quantiles by filtering all the RBP pairs with the integrated or STRING distance passing the threshold corresponding to the selected quantile (Figure S2E). For each quantile, we generated 120 RBP-centered modules, one module per RBP, by identifying all the RBPs passing the integrated or STRING distance threshold. Module sizes have been calculated as the number of interacting RBPs plus 1 (including the RBP of origin). For STRING, the distances were calculated as 1000 - STRING score, all 7140 RBP pairs were included in this analysis.

#### Alternative splicing events analysis

RNA-seq data was processed as follows: (1) to fulfill MISO requirements (see below), the reads obtained with different sequencing lengths were truncated to 75bps with *cutadapt* (v.2.10) -l option, (2) the truncated reads were mapped to the human hg38 genome assembly with GENCODE v38 comprehensive gene annotation using *STAR* (v.2.7.9) with options --outFilterScoreMinOverLread and --outFilterMatchNminOverLread both set to 0.25 (Dobin et al. 2013), (3) non-unique alignment were filtered and the replicates were merged, (4) the insert size distribution was estimated for each merged bam file separately using pe_utils –compute-insert-len from *MISO* (v.0.5.4), constitutive exons were retrieved using exon_utils with --get-const-exons and --min-exon-size 1000 (Katz et al. 2010), (5) alternative splicing events were identified using miso --run with --read-len set to 75 and --paired-end set to the previously estimated insert size parameters. Finally, only cases with non-zero numbers of exclusion and inclusion reads and sum of these reads >=10 in at least one sample are left and shown in Figure 4.

#### Ribosome profiling analysis

To process the reads, the Ribo-seq reads were first trimmed using *cutadapt* (v2.3) to remove the linker sequence AGATCGGAAGAGCAC. The fastx_barcode_splitter script from the *Fastx* toolkit was then used to split the samples based on their barcodes. Since the reads contain unique molecular identifiers (UMIs), they were collapsed to retain only unique reads. The UMIs were then removed from the beginning and end of each read (2 and 5 Ns, respectively) and appended to the name of each read. *Bowtie2* (v2.3.5) was then used to remove reads that map to ribosomal RNAs and tRNAs, and the retained reads were then aligned to mRNAs (we used the isoform with the longest coding sequence for each gene as the representative). Subsequent to alignment, *umitools* (v0.3.3) was used to deduplicate reads.

The quality check and downstream processing of the processed reads was performed using *Ribolog* v0.0.0.9 (Navickas et al. 2021). To distinguish stalling peaks from stochastic sequencing artifacts, we followed a multi-step procedure. We calculated P-site offsets and identified the codon at the ribosomal A-site for each RPF read using the *riboWaltz* package. A loess smoother was used to de-noise codon-wise RPF counts. The loess span parameter varied depending on the transcript length and allowed borrowing information from approximately 5 codons on either side of the A-site. We calculated an excess ratio at each codon position by dividing the loess-smoothed count by the transcript’s background translation level (median of no-zero loess-smoothed counts). After median normalization of the corrected counts and removal of transcripts with 0 counts, the ribosome occupancy testing was performed using logistic regression in *Ribolog*.

#### Perturb-seq analysis

Cell Ranger (version 3.0.1, 10X Genomics) with default parameters was used to align reads and generate digital expression matrices from single-cell sequencing data. To assign cell genotypes, a bwa reference (H. Li and Durbin 2009) database was created containing all guide sequences present in the library using bwa index command. The barcode-enrichment libraries were mapped to this database to establish the guide identities; to detect the cell barcodes, the barcode correction scheme used in Cell Ranger was used (the mapping of uncorrected to corrected barcodes was extracted from Cell Ranger analysis run of the whole transcriptome libraries; this mapping was then applied to the reads of barcode-enrichment libraries). UMI correction was performed by merging the UMIs within the hamming distance of 1 from each other. For each UMI, the guide assignment was done by choosing the guide sequence most represented among the reads containing the given UMI. To make the final assignment of a guide to cell barcodes, we only considered the barcodes that were represented by at least 5 different UMIs, with >80% UMIs representing the same guide.

Data filtering was performed using *scanpy* package (F. A. Wolf, Angerer, and Theis 2018) . Data were denoised using a modification of *scvi* autoencoder (Gayoso et al. 2022) with loss function penalizing for the similarity between cells having different RBPs knocked down. The distance between transcriptome profiles of individual RBP knockdowns was calculated by applying the t-test to individual gene counts across the cells that were assigned the respective guide sequence.

#### MS data analysis (BioID2 mass spectrometry data)

Data were quantified and queried against a Uniprot human database (January 2013) using *MaxQuant* MaxLFQ command (Cox et al. 2014). Data normalization was performed in Perseus (Tyanova et al. 2016) (version 1.6.2.1). For batch correction, Brent Pedersen’s implementation (Pedersen n.d.) of ComBat function from *sva* package (Leek et al. 2012) was used. The protein abundances in “experiment” (biotin +) and “control” (biotin -) samples were compared using t-test for each protein individually.

#### ATAC-seq analysis

ENCODE ATAC-seq pipeline (Lee et al. 2016) with default parameters was used for sequencing data processing and analysis. The differentially accessible peaks were identified with the DESeq2 package (Love, Huber, and Anders 2014) and annotated with the *ChIPseeker* package (Yu, Wang, and He 2015). To perform a comparison against published ChIP-Seq data, the processed ChIP-exo results were downloaded from GEO (GSE151287) (Lai et al. 2021) The data consisted of bed files containing 33 and 181 QKI peaks (two replicates) and a bigWig file with ZNF800 ChIP-exo signal (no ChIP-exo peaks were reported for ZNF800). In total, 234564 and 222350 ATAC-seq peaks for QKI and ZNF800, respectively, had coverage of at least 10 reads in more than one replicate and were used in the following tests. For QKI, the bed files with ChIP-exo peaks were merged, transferred to the hg38 genome assembly with UCSC *liftOver* and the numbers of differentially accessible (or not differentially accessible) QKI-KD ATAC-seq peaks that intersect (or do not intersect) ChIP-exo peaks were calculated using bedtools intersect (v.2.26.0) (Hinrichs et al. 2006; Quinlan and Hall 2010) followed by a one-sided (’greater’) Fisher’s exact test on 2x2 contingency table. For ZNF800, bigWig files were converted to bed using UCSC bigWigToWig (v.377) and wig2bed from BEDOPS (v.2.4.38) (Kent et al. 2010; Neph et al. 2012), followed by UCSC *liftOver* to the hg38 genome assembly. The resulting regions were intersected with differentially accessible and not differentially accessible ZNF800-KD ATAC-seq peaks using bedtools intersect followed by comparison of ChIP-exo signal distribution in these two sets using non-parametric Mann-Whitney U test.

#### MS data analysis (TAF15 KD proteomic quantification)

Quantitative analysis of the TMT experiments was performed simultaneously to protein identification using *Proteome Discoverer 2.5* software. The precursor and fragment ion mass tolerances were set to 10 ppm, 0.6 Da, respectively), enzyme was Trypsin with a maximum of 2 missed cleavages and Uniprot Human proteome FASTA file and common contaminant FASTA file used in SEQUEST searches. The impurity correction factors obtained from Thermo Fisher Scientific for each kit was included in the search and quantification. The following settings were used to search the data; dynamic modifications; Oxidation / +15.995Da (M), Deamidated / +0.984 Da (N, Q), Acetylation /+42.011 Da (N-terminus) and static modifications of TMT6plex / +229.163 Da (N-Terminus, K), MMTS / +45.988 Da (C).

*Scaffold Q+* (version Scaffold_5.0.1, Proteome Software Inc., Portland, OR) was used to quantitate TMT Based Quantitation peptide and protein identifications. Peptide identifications were accepted if they could be established at greater than 78.0% probability to achieve an FDR less than 1.0% by the Percolator posterior error probability calculation (Käll, Storey, and Noble 2008). Protein identifications were accepted if they could be established at greater than 5.0% probability to achieve an FDR less than 1.0% and contained at least 1 identified peptides. Protein probabilities were assigned by the Protein Prophet algorithm (Nesvizhskii et al. 2003). Proteins that contained similar peptides and could not be differentiated based on MS/MS analysis alone were grouped to satisfy the principles of parsimony. Proteins sharing significant peptide evidence were grouped into clusters. Channels were corrected by the matrix [0.000,0.000,0.931,0.0689,0.000]; [0.000,0.000,0.933,0.0672,0.000]; [0.000,0.00750,0.931,0.0619,0.000]; [0.000,0.0113,0.929,0.0593,0.000]; [0.000,0.0121,0.934,0.0532,0.000934]; [0.000,0.0148,0.923,0.0499,0.0120]; [0.000,0.0251,0.931,0.0438,0.000]; [0.000,0.0206,0.936,0.0431,0.000]; [0.000,0.0291,0.937,0.0337,0.000]; [0.000,0.0776,0.892,0.0303,0.000] in all samples according to the algorithm described in i-Tracker (Shadforth et al. 2005). Normalization was performed iteratively (across samples and spectra) on intensities, as described in Statistical Analysis of Relative Labeled Mass Spectrometry Data from Complex Samples Using ANOVA (Oberg et al. 2008). Means were used for averaging. Spectra data were log-transformed, pruned of those matched to multiple proteins, and weighted by an adaptive intensity weighting algorithm. Of 22889 spectra in the experiment at the given thresholds, 20372 (89%) were included in quantitation. Differentially expressed proteins were determined by applying T-Test with unadjusted significance level p < 0.05 corrected by Benjamini-Hochberg.

