## Supplementary Figures for "Systematic Identification of Post-Transcriptional Regulatory Modules"

**The PDF file includes:**

Figure S1: Multi-modal integration of RBP interaction data
Figure S2. RBPs interaction heatmap inferred from individual data modalities
Figure S3. BioID2-mediated proximity protein labeling
Figure S4. TAF15 and ZC3H11A regulate alternative splicing
Figure S5. TAF15 and ZC3H11A control mRNA translation and stability
Figure S6. ZNF800 and QKI control gene expression at transcriptional and post-transcriptional level

**Other Supplementary Material for this manuscript includes the following:**

Data file S1 (Microsoft Excel format). Integrated RBP Regulatory Map: pairwise distances between RNA binding proteins
Data file S2 (Microsoft Excel format). The pairwise distances between RBPs inferred from STRING-DB
Data file S3 (Microsoft Excel format). The pairwise distances between RBPs inferred from BioID2 data
Data file S4 (Microsoft Excel format). The pairwise distances between RBPs inferred from eCLIP data
Data file S5 (Microsoft Excel format). The pairwise distances between RBPs inferred from Pertub-seq data
Data file S6 (Microsoft Excel format). BioID2 Proximity Interactome: log fold changes
Data file S7 (Microsoft Excel format). BioID2 Proximity Interactome: P-values
Data file S8 (Microsoft Excel format). eCLIP RBP-RNA binding profiles
Data file S9 (Microsoft Excel format). Pathway enrichment for RNA binding proteins inferred from BioID2 data: GSEA NES scores, GO Biological Process annotations
Data file S10 (Microsoft Excel format). Pathway enrichment for RNA binding proteins inferred from BioID2 data: GSEA NES scores, GO Molecular Function annotations
Data file S11 (Microsoft Excel format). Pathway enrichment for RNA binding proteins inferred from BioID2 data: GSEA NES scores, GO Cellular Component annotations
Data file S12 (Microsoft Excel format). Pathway enrichment for RNA binding proteins inferred from BioID2 data: GSEA adjusted p-values, GO Biological Process annotations
Data file S13 (Microsoft Excel format). Pathway enrichment for RNA binding proteins inferred from BioID2 data: GSEA adjusted p-values, GO Molecular Function annotations
Data file S14 (Microsoft Excel format). Pathway enrichment for RNA binding proteins inferred from BioID2 data: GSEA adjusted p-values, GO Cellular Component annotations
Data file S15 (Microsoft Excel format). Data availability for the studied RNA binding proteins

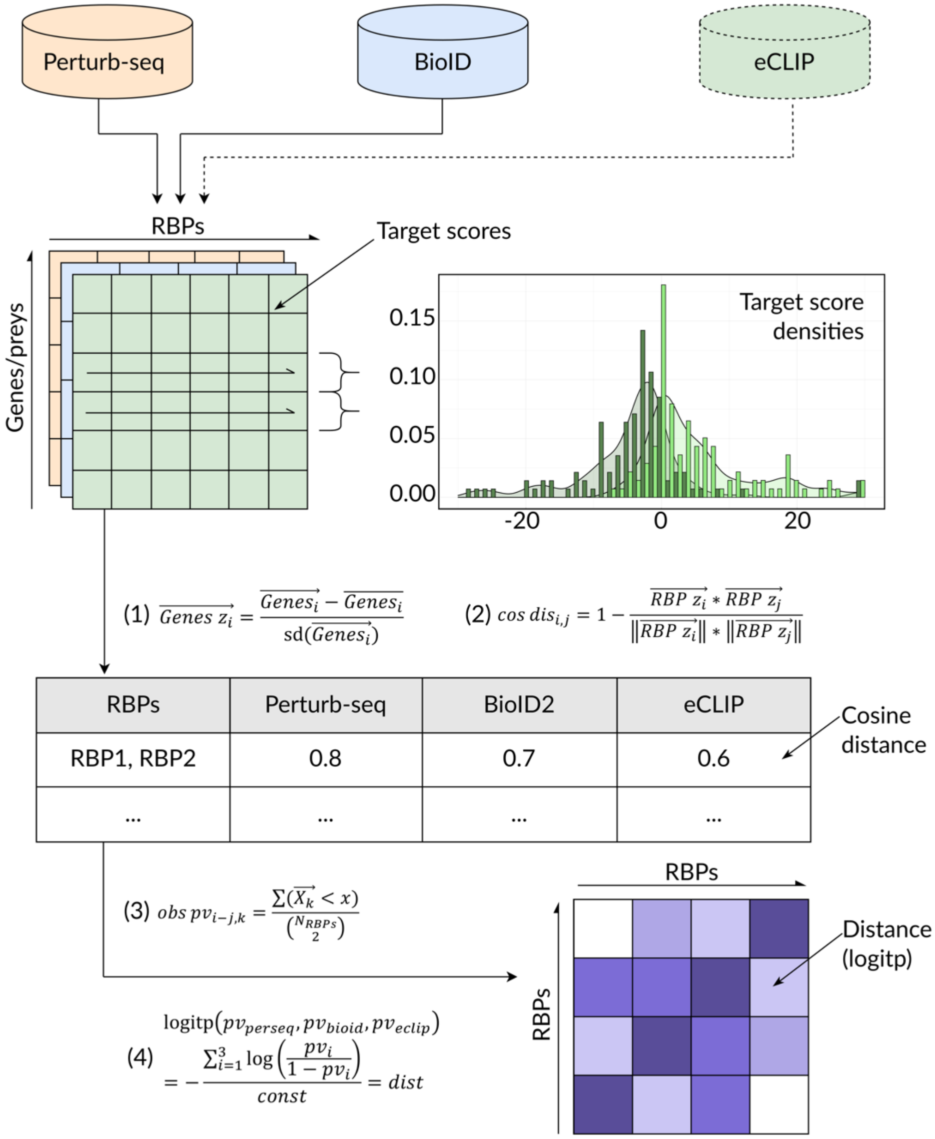


**Figure S1. Multi-modal integration of RBP interaction data**

Top: the three data modalities are shown on top; the datasets generated in this paper are highlighted with solid line, and the data downloaded from publicly available resources is highlighted with dashed line.

Middle: each dataset was preprocessed into a table, where the columns are RBPs and the rows are gene targets (shown in color for individual datasets). Every RBP was represented by a numeric column vector. The gene targets correspond to: for Perturb-Seq - individual genes, for BioID2 - protein binding partners, for eCLIP - mRNA binding targets. Difference between two numeric row vectors is shown on the right in the form of a histogram.

Bottom: the formulas applied at the key steps of the integration procedure are shown. (1): numeric column vectors were normalized by applying z-score transformation. (2): For each dataset, the cosine distances between pairs of individual RBPs were calculated. (3): The resulting distances were then transformed into empirical P-values reflecting assay-specific inter-RBPs distances. (4): Finally, a single interaction score was measured for each RBP pair by combining the P-values from the three assays using logit aggregation.

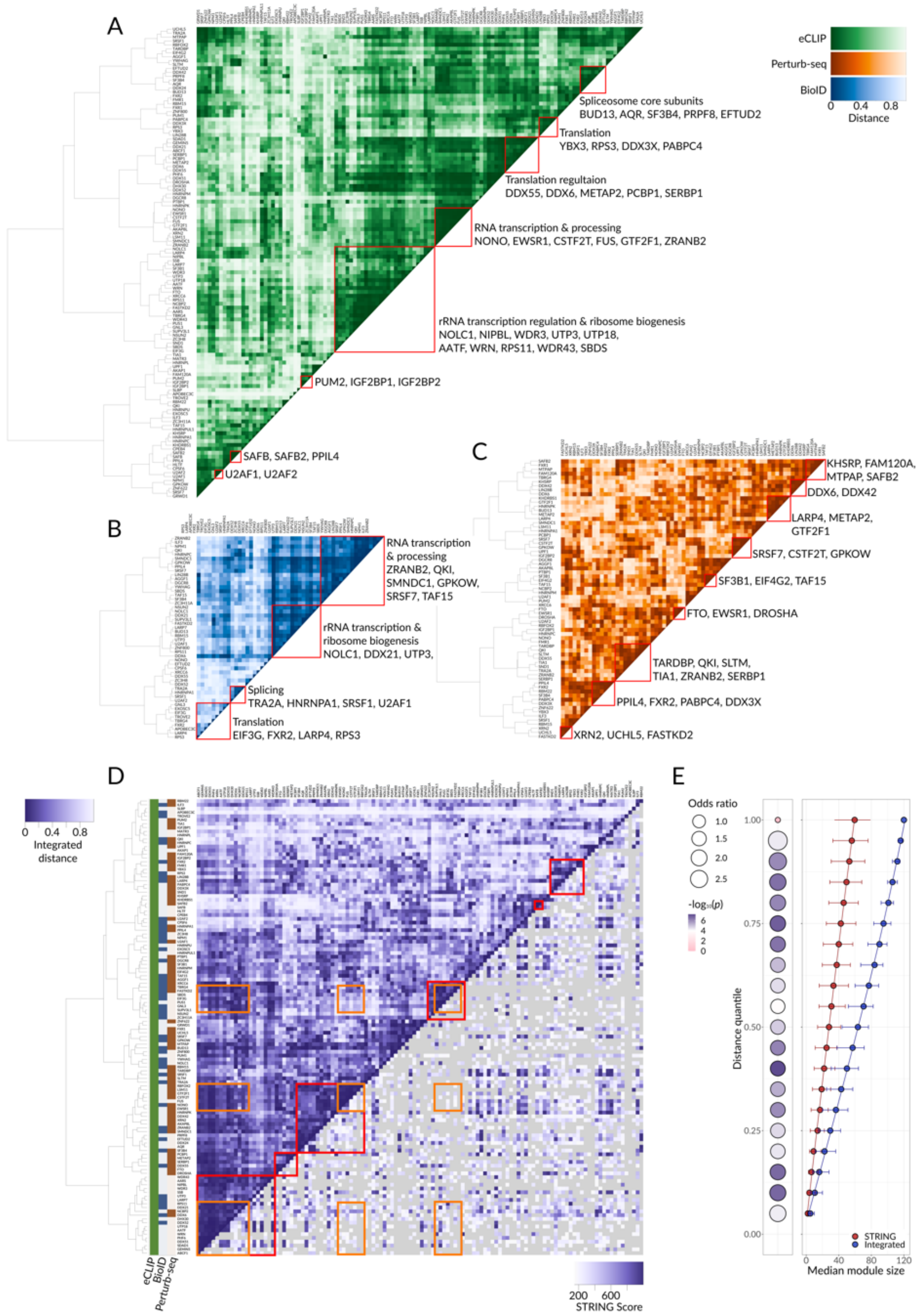


**Figure S2. RBPs interaction heatmap inferred from individual data modalities**

**(A)** A heatmap showing the pairwise distances between RBPs as informed by eCLIP data. Each row and column represents a given RBP; the hierarchical clustering dendrogram of RBPs is shown on the left. Known regulatory modules, consisting of previously annotated functional interactions, are marked with red borders and labeled.

**(B)** A heatmap showing the pairwise distances between RBPs as informed by the BioID2 dataset. Annotations as in (A).

**(C)** A heatmap showing the pairwise distances between RBPs as informed by Perturb-seq dataset. Annotations as in (A).
**(D)** Upper triangle: the heatmap of Integrated RBP Regulatory Map as in Fig. 2A. Lower triangle: the heatmap showing the pairwise distances between RBPs inferred from STRING-DB (Szklarczyk et al. 2018). The location of RBPs within the heatmap is a symmetrical reflection of the upper triangle. The pairs of RBPs where the interaction score could not be calculated are shown in gray. The same regulatory modules as in Fig. 2A are highlighted in red and yellow.
**(E)** Left subpanel: intersection between the interactions in STRING database (STRING score >= 700) and those in the 'integrated' distance modules passing particular quantiles of the distance distribution. 3403 RBP pairs with the defined STRING score (of 7140 total pairs) were used as the superset to compute the p-value of the overlap (one-sided Fisher's exact test). Right subpanel: median sizes of the RBP-centered modules assembled using integrated distance or STRING score-based distance. All 7140 RBP pairs were used, error bars correspond to lower and upper quartiles.

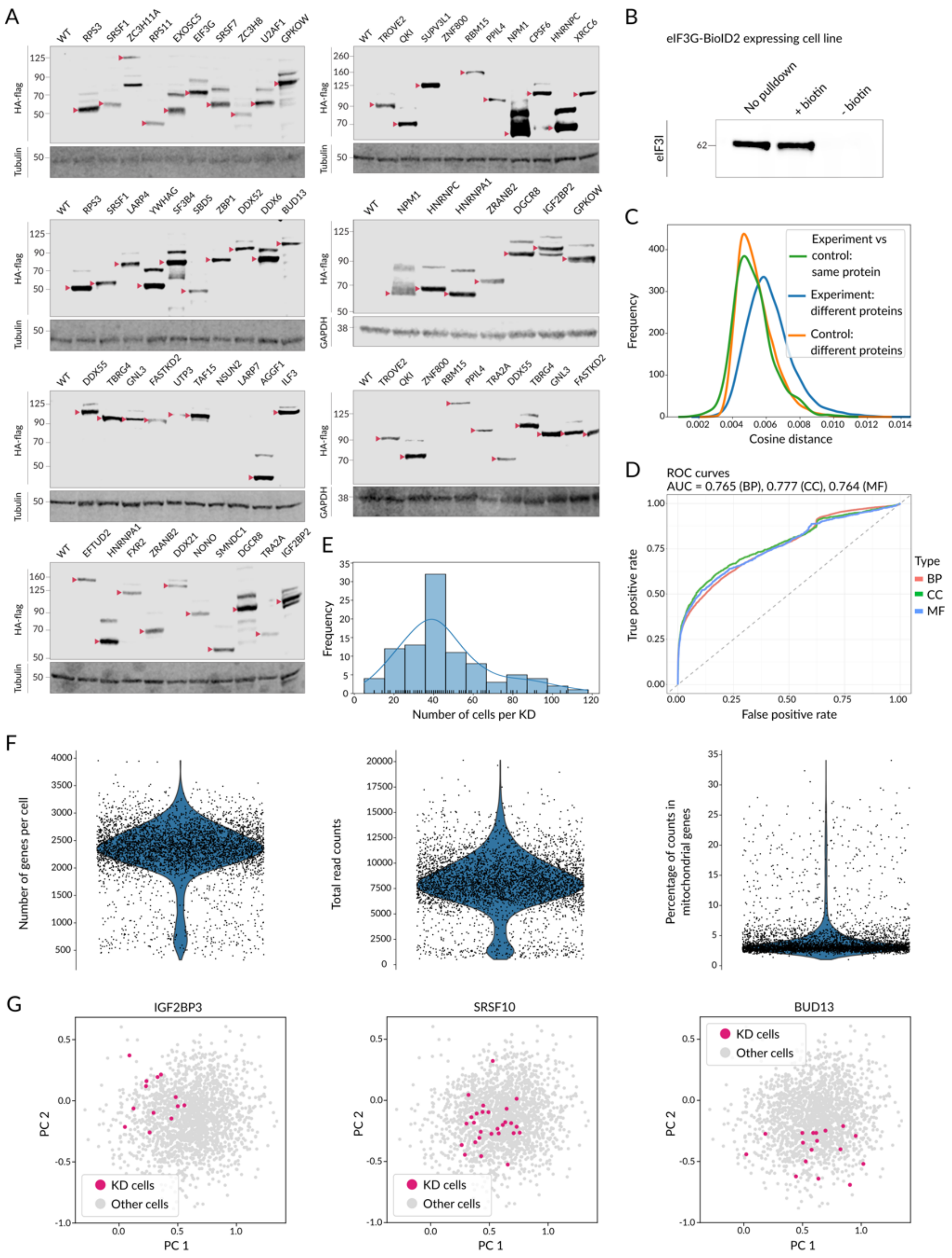


**Figure S3. BioID2-mediated proximity protein labeling**

**(A)** Western blot analysis of cell lysates collected from each of 50 RBP-BioID2-expressing cell lines. HA antibody along with either tubulin or GAPDH antibodies for endogenous controls were used. Full blots are shown, the lanes are labeled by the fusion protein expressed; the bands corresponding to the correct protein sizes are highlighted by red arrows.

**(B)** Western blot analysis of eIF3I in the lysate collected from eIF3G-BioID2-expressing cell line. Input lysate, streptavidin pulldown sample, and negative control pulldown (without the addition of biotin) are shown.
**(C)** Kernel Distribution Estimation Plot of pairwise cosine distances between proteomic profiles of all the individual samples. The pairwise distances were grouped into categories based on two features: (1) if the samples are “experiments” (+ biotin) or “control” (- biotin), (2) if the samples come from the same cell line or two different cell lines. The distributions of pairwise distances within 3 representative categories are shown in color.
**(D)** The ROC curves for predictors of gene ontology (GO) annotations from RBP protein neighborhoods. BioID2-based proximity labeling data was used to predict GO annotations of RBPs as described in Methods. Then, the known GO annotations assigned to a given RBP were used to estimate the specificity and sensitivity of the classifier. The curves corresponding to 3 types of GO annotations are shown in color: BP, MF, and CC represent Biological Process, Molecular Function, and Cellular Component groups of GO, respectively.
**(E)** A histogram and a Kernel Distribution Estimation Plot showing the number of single cell transcriptomes sequenced for each individual RBP knockdown for the Perturb-seq dataset.
**(F)** Violin plots showing the number of genes sequenced, number of total reads, and percentage of reads mapping to mitochondrial genes for the single cell transcriptomes sequenced. The cells whose corresponding sgRNA was successfully identified are shown.
**(G)** PCA projections of the single cell transcriptomes. The cells carrying knock downs of a target RBP are highlighted in red. The knockdown cells are highlighted for example proteins IGF2BP3 (left), SRSF10 (middle) and BUD13 (right).

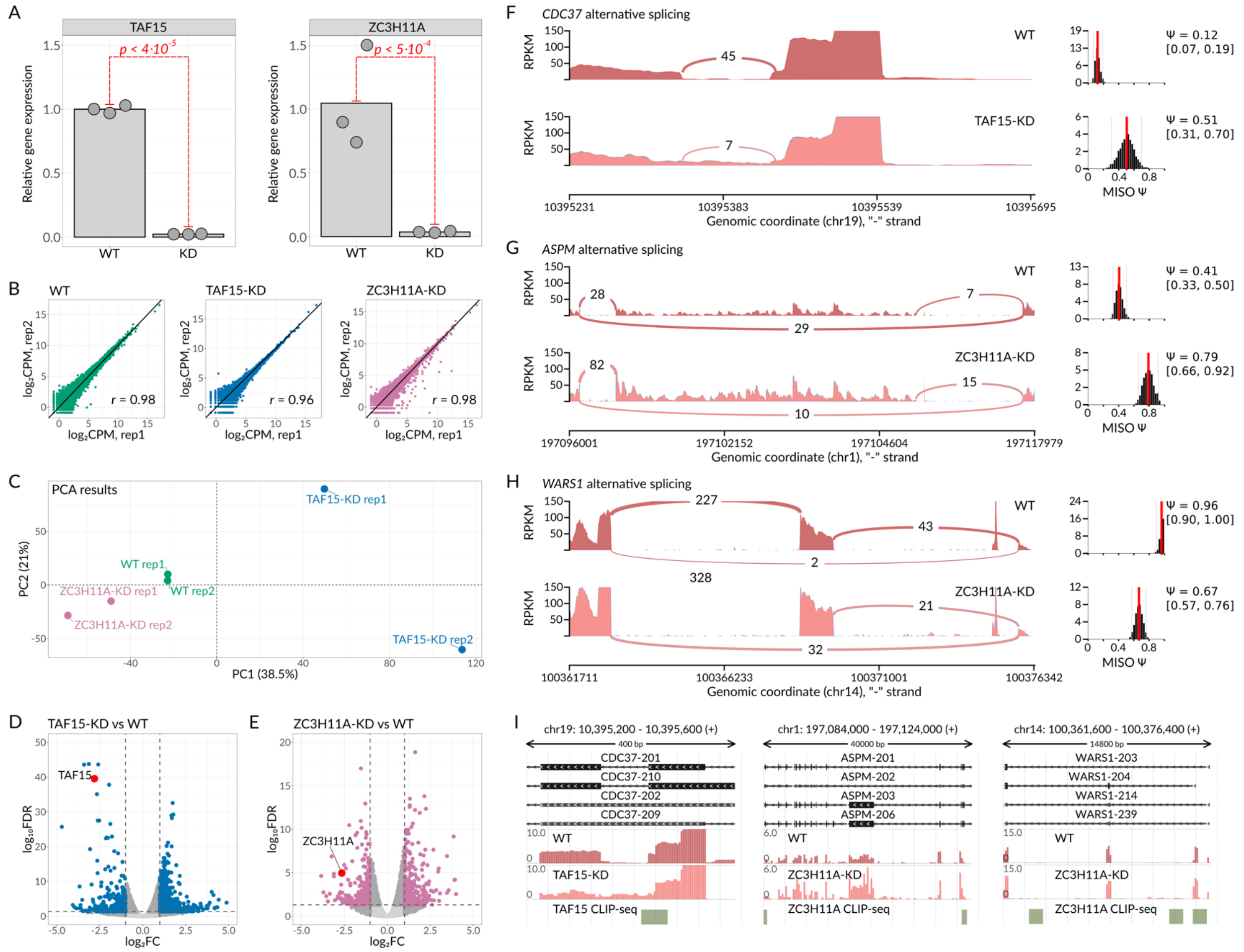


**Figure S4. TAF15 and ZC3H11A regulate alternative splicing**

**(A)** RT-qPCR quantification of relative levels of *TAF15* and *ZC3H11A* transcripts in the respective knockdown cell lines. *P* from t-test performed on log-transformed expression estimates.

**(B)** Scatter plots showing the correlations between biological replicates log2CPM for RNA-seq experiments.

**(C)** PCA analysis of RNA-seq samples for WT, TAF15-KD and ZC3H11A-KD cells. The first two principal components are shown. Analysis was performed using log2CPM values.

**(D)** Volcano plot showing the changes in gene expression upon TAF15 knockdown. *TAF15* is highlighted in red. The vertical dashed lines show log2FC thresholds of -1 and 1. The horizontal dashed line corresponds to FDR of 0.05. In total, there are 930 genes passing |log2FC| > 1 and FDR < 0.05 (highlighted in blue)

**(E)** Volcano plot showing the changes in gene expression upon ZC3H11A knockdown with 565 genes passing |log2FC| > 1 and FDR < 0.05 filters (highlighted in pink). The filters are shown as in (D). *ZC3H11A* is highlighted in red.

**(F)** Sashimi plot illustrating the changes in intron retention event usage in *CDC37* transcript upon TAF15 knockdown.

**(G)** Sashimi plot illustrating the changes in skipped exon usage in *ASPM* transcript upon ZC3H11A knockdown.

**(H)** Sashimi plot illustrating the changes in skipped exon usage in *WARS1* transcript upon ZC3H11A knockdown.

**(I)** Genomic views of the *CDC37* retained intron (left), *ASPM* skipped exon (middle), and *WARS1* skipped exon (right). Below, the RNA-seq profiles from WT, TAF15-KD and ZC3H11A-KD cells are shown. Y axis: counts per million (CPM). TAF15 and ZC3H11A CLIP-seq peaks are shown at the bottom.

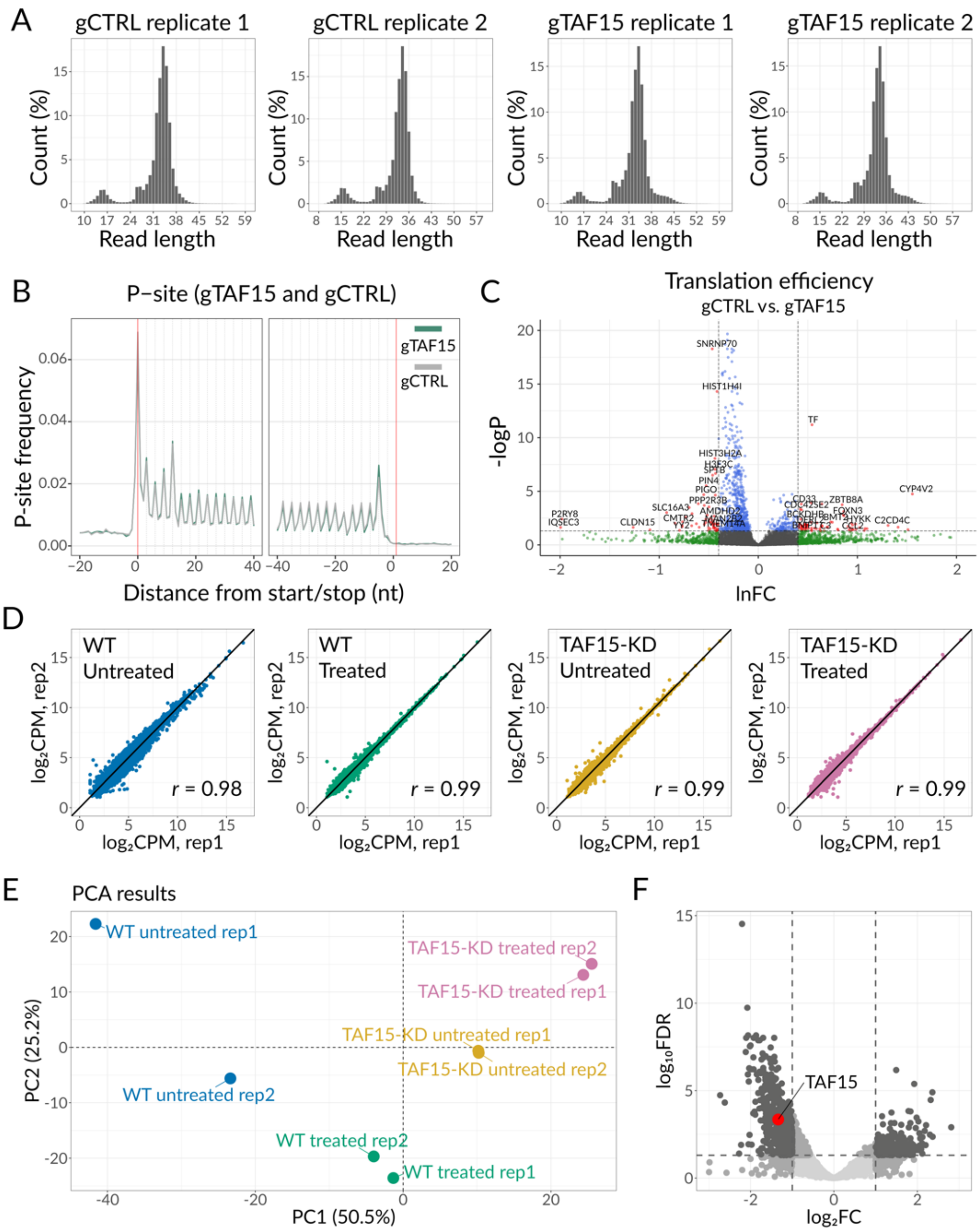


**Figure S5. TAF15 and ZC3H11A control mRNA translation and stability**

**(A)** Distribution of RPFs, aligned on an inferred ribosome P-site, on a metagene, centered around the translation start (left) or stop (right) site, for Ribo-seq of TAF15-KD and WT cells.

**(B)** Length distribution of ribosome protected footprints (RPFs) as determined by Ribo-seq.

**(C)** Volcano plot illustrating the changes in ribosome occupancy in TAF15-KD compared to control K562 cells, as determined by ribosome profiling analysis. The data points are colored according to thresholds in effect size (lnFC ± ln1.5) and significance (p-adjusted < 0.05, t-test). The genes passing both significance filters are labeled.

**(D)** Scatter plots showing the correlations between biological replicates log2CPM for RNA-seq analysis of α-amanitin treated and untreated WT and TAF15-KD cells. The different treatments and cell lines are shown in color.

**(E)** PCA analysis of RNA-seq samples for α-amanitin treated and untreated WT and TAF15-KD cells. The first two principal components are shown. Analysis was performed using log2CPM values.
**(F)** Volcano plot showing genes differential stability upon α-amanitin treatment, in TAF15-KD compared to control K562 cells. *TAF15* is highlighted red. The vertical dashed lines show log2FC thresholds of -1 and 1. The horizontal dashed line corresponds to FDR of 0.05. In total, there are 1015 genes passing |log2FC| > 1 and FDR < 0.05 (highlighted in dark gray).

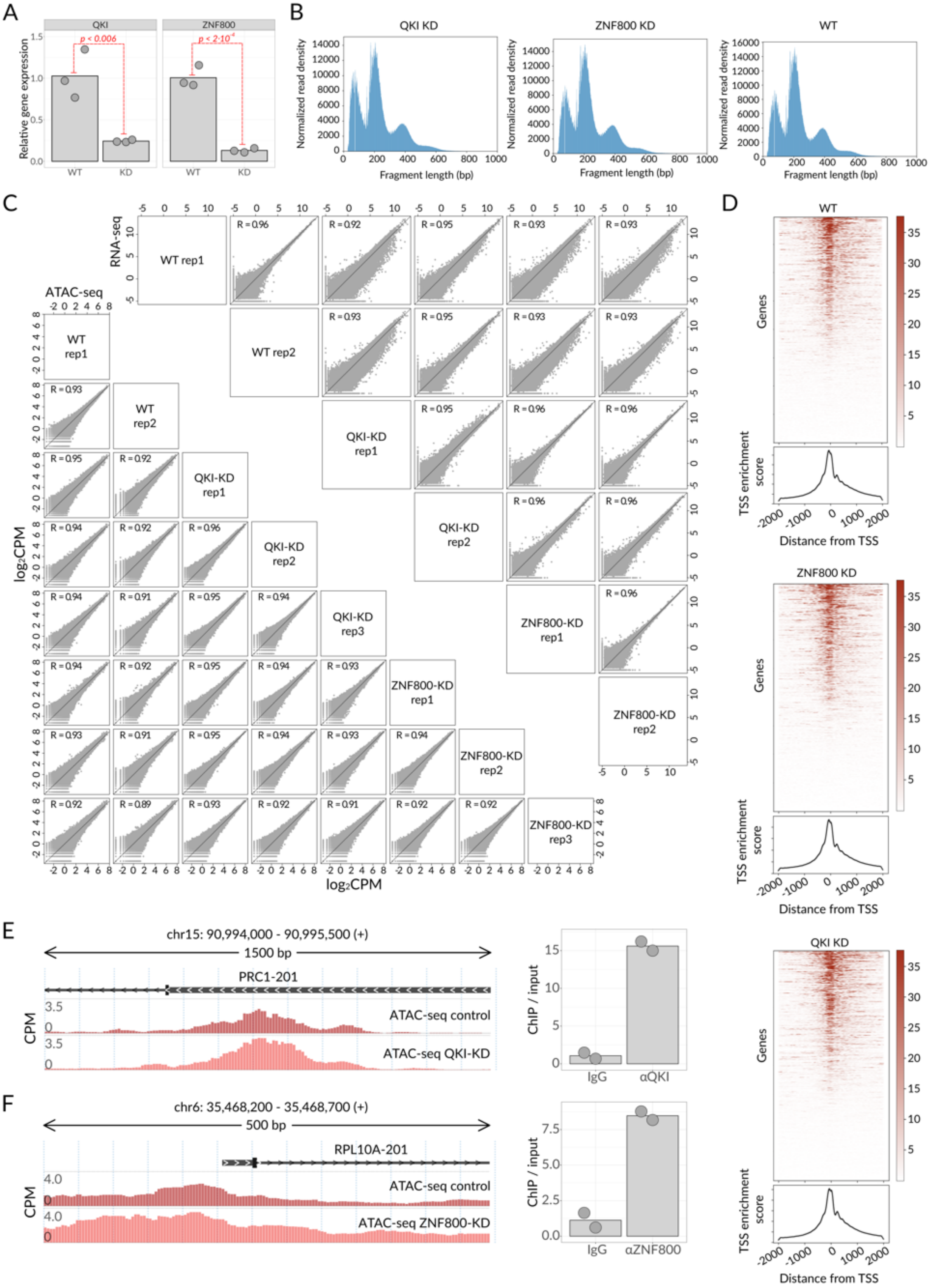
A
QKI
ZNF800
p < 0.006
$p < 2\cdot10^{-4}$
Relative gene expression
WT
KD
B
QKI KD
ZNF800 KD
WT
Normalized read density
Fragment length (bp)
C
RNA-seq
ATAC-seq
WT rep1
WT rep2
QKI-KD rep1
QKI-KD rep2
QKI-KD rep3
ZNF800-KD rep1
ZNF800-KD rep2
ZNF800-KD rep3
log₂CPM
D
WT
ZNF800 KD
QKI KD
Genes
TSS enrichment score
Distance from TSS
E
chr15: 90,994,000 - 90,995,500 (+)
1500 bp
PRC1-201
CPM
ATAC-seq control
ATAC-seq QKI-KD
ChIP / input
IgG
αQKI
F
chr6: 35,468,200 - 35,468,700 (+)
500 bp
RPL10A-201
ATAC-seq control
ATAC-seq ZNF800-KD
IgG
αZNF800

**Figure S6. ZNF800 and QKI control gene expression at transcriptional and post-transcriptional level**

**(A)** RT-qPCR quantification of relative levels of *QKI* and *ZNF800* transcripts in the respective knockdown cell lines. *P* from t-test performed on log-transformed expression estimates.

**(B)** Histograms showing the fragment length distributions for ATAC-seq experiments for QKI-KD cells (left), ZNF800-KD cells (middle), and WT cells (right). A single exemplar replicate is shown per experiment.

**(C)** Scatter plots showing the correlations between biological replicates log2CPM for RNA-seq experiments (top) and ATAC-seq experiments (bottom).

**(D)** Enrichment of ATAC-seq reads near transcription start sites (TSS) is shown with heatmaps and Kernel Distribution Estimation (KDE) plots for WT cells (top), ZNF800-KD cells (middle), and QKI-KD cells (bottom). Heatmaps: rows correspond to genes; columns correspond to genomic positions relative to TSSs. Cell values show the number of ATAC-seq reads aligned to a given region. KDE plots show TSS enrichment scores across genes. A single exemplar replicate is shown per experiment.

**(E)** Left: genomic view of *PRC1* promoter region. ATAC-seq profiles of WT cells and QKI-KD cells are shown. Right: binding of QKI to the *PRC1* promoter region as measured by ChIP-qPCR in K562 cells.

**(F)** Left: genomic view of *RPL10A* promoter region. ATAC-seq profiles of WT cells and ZNF800-KD cells are shown. Right: binding of ZNF800 to the *RPL10A* promoter region as measured by ChIP-qPCR in K562 cells.
